# A bovine-derived influenza A virus (H5N1) shows efficient replication in well-differentiated human nasal epithelial cells at the temperature of the upper airways

**DOI:** 10.64898/2026.01.16.699876

**Authors:** Étori Aguiar Moreira, Samuel Constant, Charlène Constant, Lisa Butticaz, Michele Wyler, Teodora David, Peter M. Grin, Charaf Benarafa, Volker Thiel, Marco P. Alves, Gert Zimmer

## Abstract

Highly pathogenic avian influenza (H5N1) viruses of clade 2.3.4.4b have caused significant losses among bird populations worldwide and have repeatedly crossed the species barrier, infecting mammals, including humans. However, efficient human-to-human transmission has not yet been observed. Here, we demonstrate that an H5N1 virus isolated from bovine milk in Texas in 2024 (H5N1_Tex/24_) replicates in differentiated human nasal epithelial cells as efficiently as a 2009 pandemic H1N1 virus strain (H1N1_HH4/09_) at both 37 °C and 33 °C. The adaptive mutations PB2-M631L and PA-K497R do not affect H5N1_Tex/24_ replication at 37 °C but promote replication at 33 °C. Conversely, H5N1_BE/22_, a virus from the same clade isolated from a pelican in 2022 that lacks these mutations, replicates in human nasal epithelial cells at 37 °C as efficiently as H5N1_Tex/24_ but exhibits limited replication at 33 °C. Introducing the two mutations PB2-M631L and PA-K497R did not overcome this limitation. Furthermore, nasal epithelial cells express receptors for both human and avian influenza viruses. Accordingly, no mutations were detected in HA which are known to switch receptor preference. Finally, we demonstrate that H5N1_Tex/24_ remains sensitive to the antiviral effects of interferon-λ (IFN-λ), however, infected nasal epithelial cells secrete only small amounts of this cytokine. Overall, our results suggest that H5N1_Tex/24_ possesses intrinsic traits enabling efficient replication in the human upper airways.

## Introduction

In recent years, H5N1 highly pathogenic avian influenza (HPAI) viruses of the phylogenetic clade 2.3.4.4b have spread worldwide, causing the death of millions of wild birds and domestic poultry^1,2^. In addition, viruses of this clade were able to infect various mammalian species including harbor seals, sea lions, foxes, minks, and cats, indicating a significantly expanded host range compared to older clades^3^. In some mammalian species such as cats and foxes, H5N1 viruses of clade 2.3.4.4b showed a marked neurotropism causing a mostly fatal disease^4,5^. In addition, clade 2.3.4.4b viruses of genotype B3.13 and genotype D1.1 were able to infect dairy cows in which they replicated in mammary gland tissue, resulting in high titers of infectious virus shed into milk^6^. This “bovine H5N1” virus was further disseminated by milking equipment and animal transport to 20 US states with about 1093 herds being affected as of March 2026^7^.

Several people who had been in close contact with infected cows or poultry were infected by H5N1 viruses of clade 2.3.4.4b^8^. Most affected persons displayed only mild symptoms such as conjunctivitis^9^, however, one adult person with underlying comorbidities died following infection^8^. In addition, a 13-year-old girl in Canada with a history of mild asthma and elevated body-mass index succumbed to infection by H5N1 of genotype D1.1^10^. No human-to-human transmission of bovine H5N1 virus has been observed to date. Likewise, airborne transmission of bovine H5N1 virus in the ferret model is lacking or inefficient^11-15^. Nevertheless, infection of ferrets with other influenza viruses suggest a strong correlation between viral loads in the upper respiratory tract, virus shedding and airborne transmission^15^.

^16^An important species barrier for efficient replication of avian influenza viruses in the human respiratory tract is the specific binding of hemagglutinin (HA) to sialylated glycoconjugates on the host cell^17,18^. Avian influenza viruses are known to preferentially bind to α2,3-linked sialic acid residues, which are less abundantly expressed in the human upper respiratory tract than α2,6-linked sialic acid, the receptor determinant for human influenza viruses^19^. There are conflicting reports on the receptor specificity of bovine-derived H5N1 viruses. On the one hand, bovine H5N1 viruses were shown to recognize both α2,3 and α2,6-linked sialic acid^11^. On the other hand, there is evidence that bovine H5N1 has retained its preference for α2,3-linked sialic acid residues^20,21^, although a single amino acid substitution in HA is sufficient to switch the preference to α2,6-linked sialic acid, the human-type receptor^22,23^. In consistence with the avian-type receptor usage and the lower prevalence of α2,3-linked sialic acids in the human upper respiratory tract, bovine H5N1 virus showed inefficient binding to these tissues and replicated more inefficiently in nasal epithelial cell cultures compared to a seasonal H3N2 virus^24^. Previous gain-of-function experiments in ferrets suggested that adaptation to α2,6-linked sialic acids contributes to both infection of cells in the human nasal conchae and airborne virus transmission^25^.

The adaptation of avian influenza viruses to replication in human tissue is often associated with characteristic amino acid substitutions in the PB2 protein such as E627K^26,27^, D701N^28^, and Q591R^26,27,29^. Interestingly, the PB2-E627K substitution is present in a B3.13 H5N1 virus isolated from the first human case infected with a genotype B3.13 H5N1 virus^30^,but was virtually absent from most bovine H5N1 virus isolates^31^, although it popped up in dairy cows experimentally infected with European clade 2.3.4.4b HPAI virus^32^. Instead of E627K, the PB2 protein of bovine H5N1 viruses frequently contains the M631L substitution, which, like PB2-E627K, leads to increased activity of the viral polymerase complex in mammalian cells ^12,33-35^. The PB2-M631L substitution allows the polymerase to better interact with bovine ANP32 proteins, host factors that are essentially needed to stabilize the viral polymerase complex^36^. The H5N1 polymerase activity is further increased in mammalian cells by the PA-K497R substitution present in 95% of bovine H5N1 viruses^35^. Interestingly, neither the PB2-E627K nor the PB2-M631L substitution but the PB2-D701N is found in H5N1 viruses of genotype D1.1 that spread to dairy cows in Nevada and Arizona^37^.

Another factor that influences the replication of avian influenza viruses in the human upper respiratory tract is temperature. While the physiological temperature in the nasal cavity ranges from 25 °C (at the nares) to 33 °C (in the nasopharynx), avian influenza viruses are typically adapted to replication at 41 - 42 °C - the normal body temperature of avian species^38^. However, avian influenza viruses are certainly capable of adapting to these lower temperatures^26-28,39-43^. The ability of avian influenza viruses to replicate at low temperatures has been linked to the PB2-E627K mutation^44^. However, there is also evidence for a role of the envelope glycoproteins HA and NA in replication at low temperatures^42^.

The innate immune response also poses a significant barrier for zoonotic viruses. The interferon (IFN)-stimulated MxA protein can effectively limit the replication of avian influenza viruses in the human respiratory tract^45,46^. A recent study suggests that bovine H5N1 virus remains susceptible to the antiviral effects of human MxA^47^.

The finding that bovine H5N1 viruses do not show human-to-human transmission may be due to their inability to replicate effectively in the human upper respiratory tract, for example, because of a lack of suitable receptors, an inability to overcome the innate immune response, or because the viruses are not adapted to the lower temperatures in the upper respiratory tract. To investigate these parameters in more detail, we examined the replication of a clade 2.3.4.4b H5N1 virus isolated from bovine milk in 2024 (H5N1_Tex/24_) in well differentiated human nasal and bronchial epithelial cells cultured at the air-liquid interface, comparing its replication with that of a 2009 pandemic H1N1 virus (H1N1_HH4/09_) and avian H5N1 virus strains. Multi-step replication kinetics were assessed at 33 °C and 37 °C, and potential genomic changes arising during replication in these cells were analyzed. We also evaluated the presence of avian- and human-type influenza virus receptors in differentiated human nasal epithelial cells, as well as the induction and secretion of IFNs following infection. Our results show that H5N1_Tex/24_ replicates as efficiently as H1N1_HH4/09_ without requiring additional genetic adaptation. Notably, its robust replication in human nasal epithelial cells suggests that H5N1_Tex/24_ already possesses traits that may facilitate infection of the human upper respiratory tract, posing a potential risk to human populations lacking adequate immunity to this zoonotic virus.

## Results

### Replication of H5N1_Tex/24_ in primary human nasal epithelial cells

To assess the replication capacity of influenza A viruses in the human upper respiratory tract, we used a pseudostratified three-dimensional model of human airway epithelium (MucilAir; Epithelix) derived from primary nasal epithelial cells. Following differentiation at the air-liquid interface (ALI), basal progenitor cells develop into a polarized mucociliary epithelium composed of basal, ciliated, and mucus-producing cells, closely resembling the cellular architecture of the human nasal epithelium *in vivo*^48,49^. These cultures retain key features of functional airway epithelium, including mucus production, coordinated ciliary beating, epithelial barrier integrity, and active ion transport pathways involving CFTR, ENaC, and Na⁺/K⁺-ATPase. In addition, they respond to pro-inflammatory stimuli and secrete a broad spectrum of cytokines, chemokines, and matrix-remodeling enzymes^50,51^. Throughout this study, this model is referred to as well-differentiated human nasal epithelial (HNE) cultures.

HNE cultures were infected with A/cattle/Texas/24-029328-02/2024 (H5N1_Tex/24_) or A/Hamburg/4/2009 (H1N1_HH4/09_). H5N1_Tex/24_ was isolated in 2024 from the milk of an infected dairy cow in Texas, USA, and belongs to the phylogenetic clade 2.3.4.4b, genotype B3.13^52^. It belongs to the main clade of bovine H5N1 viruses which is characterized by the PB2-M631L and PA-K497R substitutions and accounts for about 95% of all H5N1 viruses of bovine origin^35^. H1N1_HH4/09_ was isolated from a human patient during the 2009 pandemic by swine-origin H1N1 virus^53^. This viral strain has been found to escape from restriction by human MxA through adaptive mutations in the nucleoprotein^54^, and was further characterized with respect to replication and adaptation to A549 human lung epithelial cells^55^. With exception of the P83S substitution, the H1N1_HH4/09_ HA primary sequence is identical to the corresponding HA sequence of the A/California/7/2009 (H1N1) reference strain (GenBank accession no. ACP44189). In particular, it contains a glutamine residue at position 223 (H1 numbering), which is typically found in HAs that bind to α2,6-linked sialic acid^56,57^. Here, we used reverse genetics to synthetically produce H1N1_HH4/09_ (rH1N1_HH4/09_)^58^.

HNE cultures were infected with H5N1_Tex/24_ or rH1N1_HH4/09_ using a multiplicity of infection (MOI) of 0.001 TCID_50_/cell and incubated at 37 °C for 72 h. Virus released at the apical surface was sampled at 2, 24, 48, and 72 h post infection (p.i.) and titrated on MDCK cells. rH1N1_HH4/09_ replicated efficiently in HNE cultures reaching 7.9 x 10^7^ TCID_50_/mL at 48 h p.i. (**Fig. 1a**). Replication of rH1N1_HH4/09_ occurred in the absence of exogenously added trypsin indicating that serine proteases are expressed by HNE cells that can cleave the monobasic cleavage site of rH1N1_HH4/09_ HA^59^. H5N1_Tex/24_ replicated as efficiently as rH1N1_HH4/09_, reaching a maximum titer of 10^8^ TCID_50_/mL at 48 h (**Fig. 1a**). Because the H5N1_Tex/24_ virus stock had been propagated on MDCK cells, we investigated whether adaptive mutations might have facilitated its replication in HNE cells. Viral genome sequencing revealed only a few nucleotide changes relative to the published sequence, most of which were non-synonymous (**Table 1**). Notably, a G754T mutation in segment 4 (HA-D236Y, H5 numbering) was present in 17.17% of reads in the virus stock and persisted at a lower frequency (5.82%) at 72 h p.i.. A PA-I38M substitution emerged at 72 h p.i. but was present in only 5.48% of reads (**Table 1**). We did not detect mutations associated with enhanced mammalian replication (e.g. PB2-E627K, PB2-T271A and PB2-Q591R^60^) or altered receptor specificity^22^ (**Table 2**).

**Fig. 1.**
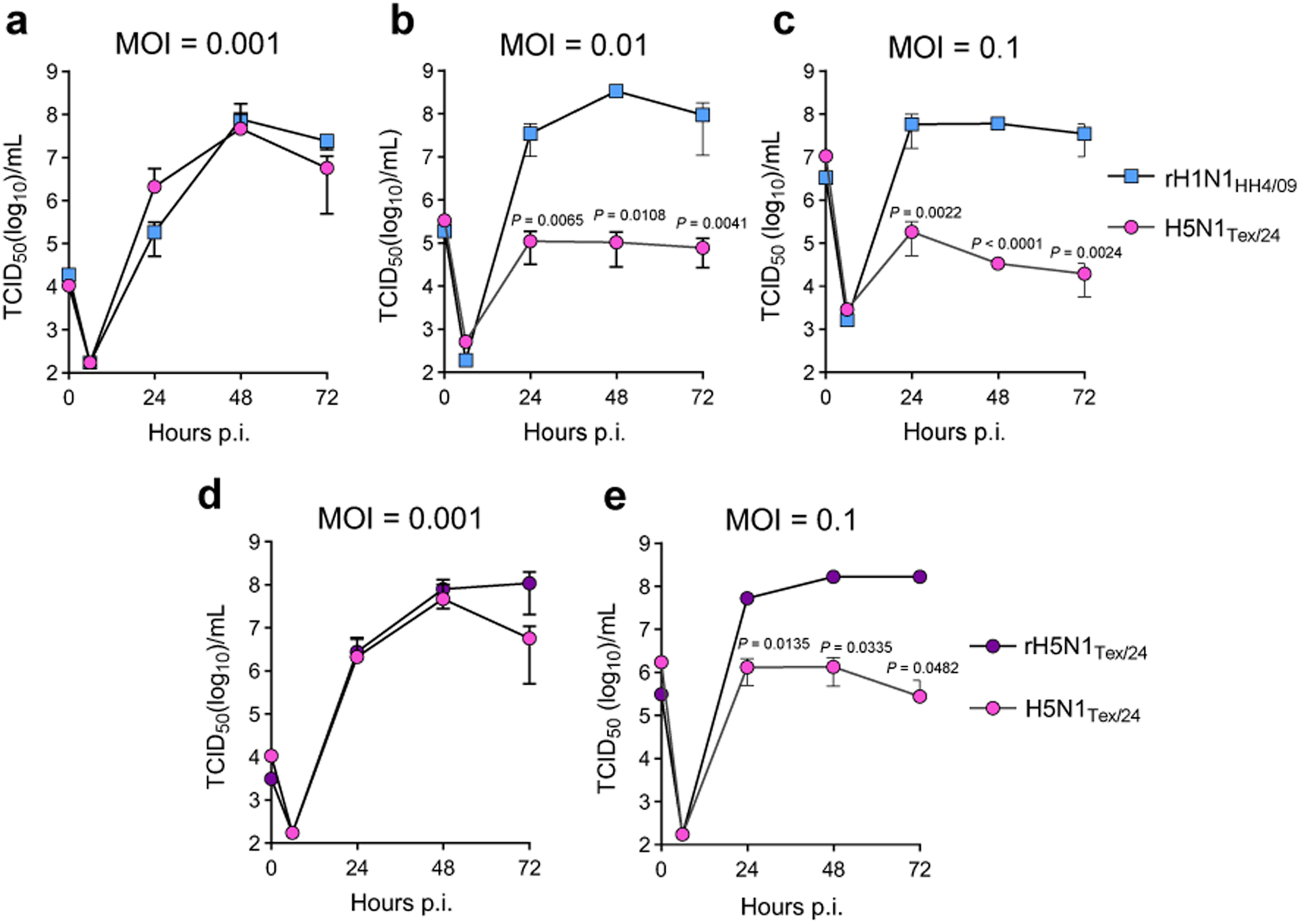
Replication kinetics of H5N1_Tex/24_ in HNE cells differentiated at the air-liquid interface. **a - c** Air-liquid interface (ALI) cultures were infected with H5N1_Tex/24_ or rH1N1_HH4/09_ at an MOI of 0.001 (**a**) 0.01 (**b**) or 0.1 (**c**). **d, e** ALI cultures were infected with H5N1_Tex/24_ or rH5N1_Tex/24_ at an MOI of 0.001 (**d**) or 0.1 (**e**). At the indicated time points, the apical surface was washed and infectious virus titers in the wash fluid were determined by endpoint dilution assay on MDCK cells. Data represent mean values ± SD from three biological replicates. Statistical significance was determined by two-way ANOVA with Sidak′s multiple comparisons test. Only statistically significant *P* values are shown in the graphs.

**Table 1.** Mutations identified in the H5N1Tex/24 genome before and after replication in HNE cells.

| Viral RNA segment | Protein | Nucleotide change<sup>a,b</sup> | Virus stock<sup>c</sup> |  | Passage in HNE<sup>d</sup> |  |
| --- | --- | --- | --- | --- | --- | --- |
|  |  |  | Freq. (%) | AA subst. | Freq. (%) | AA subst. |
| 1 | PB2 | T165C | 16.29 | - | 22.31 | - |
|  |  | G525A | 20.18 | - | 35.89 | - |
| 2 | PB1 | - | - | - | - | - |
| 3 | PA | A114G | - | - | 5.48 | I38M |
|  |  | T1545C | 19.97 | - | 5.68 | - |
| 4 | HA | G754T | 17.17 | D236Y | 5.82 | D236Y |
| 5 | NP | A1140G | 22.75 | - | 33.9 | - |
| 6 | NA | A1305C | 20.47 | - | 19.69 | - |
| 7 | M1, M2 | - | - | - | - | - |
| 8 | NS1, NEP | - | - | - | - | - |

<sup>a</sup> The sequence of the passaged virus was compared with the sequence of A/cattle/Texas/063224-24-1/2024 (H5N1) (GISAID accession number: EPI\_ISL\_19155861)
<sup>b</sup> The nucleotide position is based on the start codon.
<sup>c</sup> For stock production H5N1<sub>Tex/24</sub> was propagated for two passages on MDCK cells.
<sup>d</sup> HNE cultures were infected at an MOI of 0.001 TCID<sub>50</sub> units per cell. Apical washes were collected at 72 h p.i. and viral genome sequences were determined.

**Table 2.**
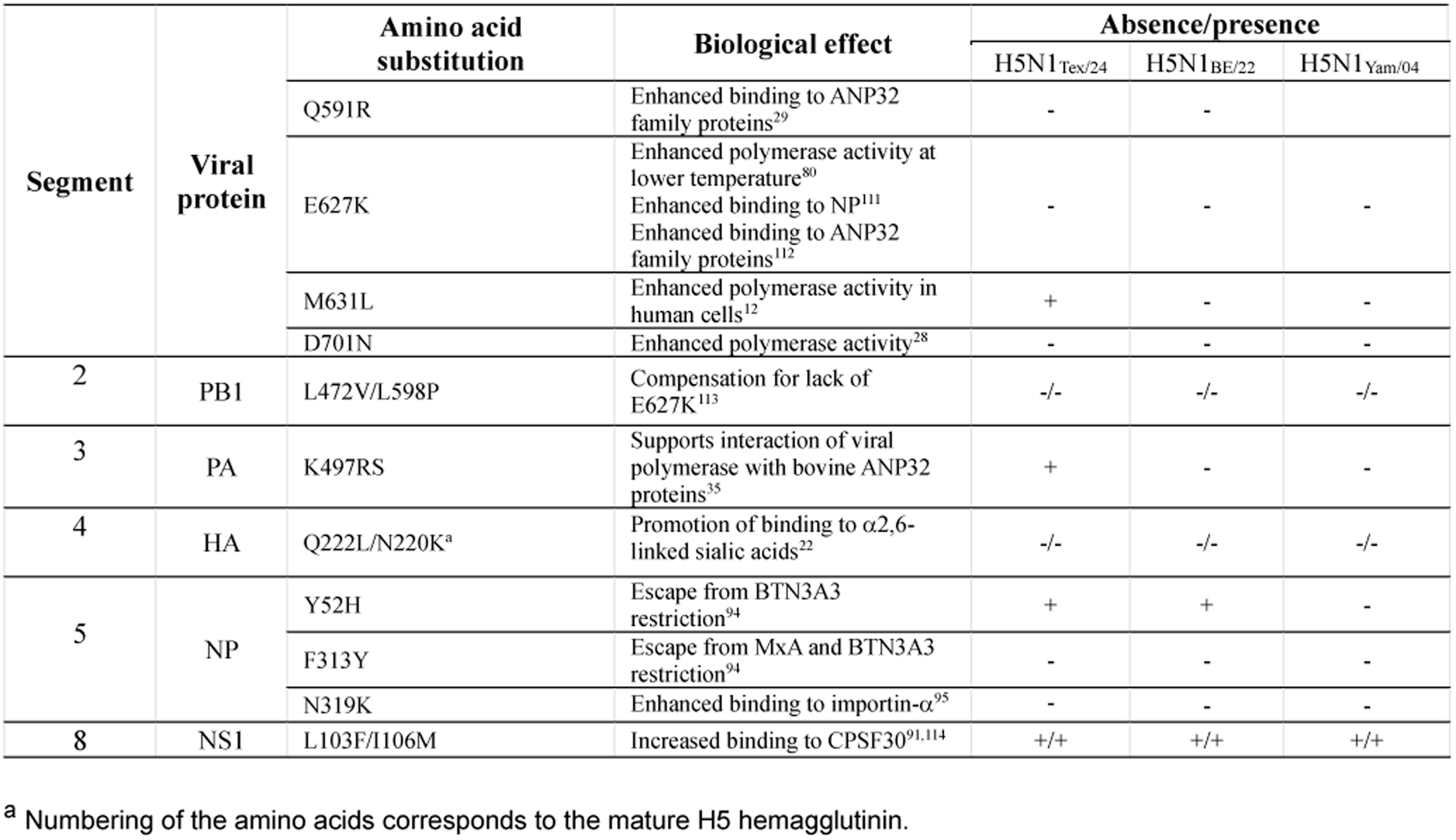
Selected mutations in H5N1 viruses adapted to mammalian hosts.

At higher inoculation doses (MOI of 0.01 or 0.1), rH1N1_HH4/09_ replicated efficiently in HNE cultures, reaching titers comparable to those observed at an MOI of 0.001 (**Fig. 1b, c**). In contrast, H5N1_Tex/24_ replication was significantly reduced compared to rH1N1_HH4/09_ if MOIs of 0.01 and 0.1 were used (**Fig. 1b, c**). To exclude effects by defective-interfering particles that might be present in the original virus isolate, we generated a recombinant H5N1 virus (rH5N1_Tex/24_) based on the published sequence of A/bovine/Texas/24-029328-02/2024 (rH5N1_Tex/24_)^61^. When HNE cells were infected using an MOI of 0.001, rH5N1_Tex/24_ replicated with kinetics comparable to the original virus isolate (**Fig. 1d**). However, if an MOI of 0.1 was used, the recombinant virus replicated more efficiently than the original virus isolate (**Fig. 1e**). These findings indicate that H5N1_Tex/24_ efficiently replicates in primary human nasal epithelial cells depending on the virus dose used, most likely because of the presence of defective-interfering particles.

### Replication of H5N1_Tex/24_ in HNE cultures at 33 °C

The human upper respiratory tract is exposed to temperatures lower than the core body temperature of 37 °C that predominates in the lower respiratory tract. To evaluate the impact of temperature on influenza A virus replication, differentiated HNE cultures were infected with rH1N1_HH4/09_, H5N1_Tex/24_, and two additional HPAI H5N1 viruses. H5N1_BE/22_, like H5N1_Tex/24_, belongs to clade 2.3.4.4b and was isolated in 2022 from a diseased Dalmatian pelican at Bern Animal Park, Switzerland. rH5N1_Yam/04_ is a recombinant clade 2.5 virus derived from a strain isolated in 2004 from infected chickens in Japan.

At 37 °C, rH1N1_HH4/09_ replicated efficiently in HNE cultures, reaching peak titers at 48 h p.i. (2.8 x 10^8^ TCID_50_/mL) before declining approximately 20-fold by 72 h p.i.. At 33 °C, replication kinetics of rH1N1_HH4/09_ were modestly delayed with peak titers observed by 72 h p.i. (**Fig. 2a**). H5N1_Tex/24_ exhibited marked attenuation during early replication at 33 °C, with infectious titers reduced by approximately 104-fold and 47-fold at 24 h and 48 h p.i., respectively, compared with 37 °C, but reached titers comparable to those at 37 °C by 72 h p.i. (**Fig. 2b**). Replication of H5N1_BE/22_ was more severely impaired at 33 °C, with titers reduced by up to 3704-fold at 72 h p.i. (**Fig. 2c**). In contrast, rH5N1_Yam/04_ replicated poorly at both temperatures (**Fig. 2d**). Analysis of infection rates further indicated significantly lower infectivity of rH5N1_Yam/04_ in HNE ALI cultures cells compared with the other viruses, despite the use of an MOI of 1 for all infections (**Supplementary Fig. 1**). Whereas H5N1_Tex/24_ and H5N1_BE/22_ replicated to levels comparable to rH1N1_HH4/09_ at 37 °C (**Fig. 2e**), only H5N1_Tex/24_ maintained similar replication efficiency 33 °C (**Fig. 2f**). Collectively, these findings suggest that H5N1_Tex/24_ displays enhanced adaptation to temperatures characteristics of the upper respiratory tract.

**Fig. 2.**
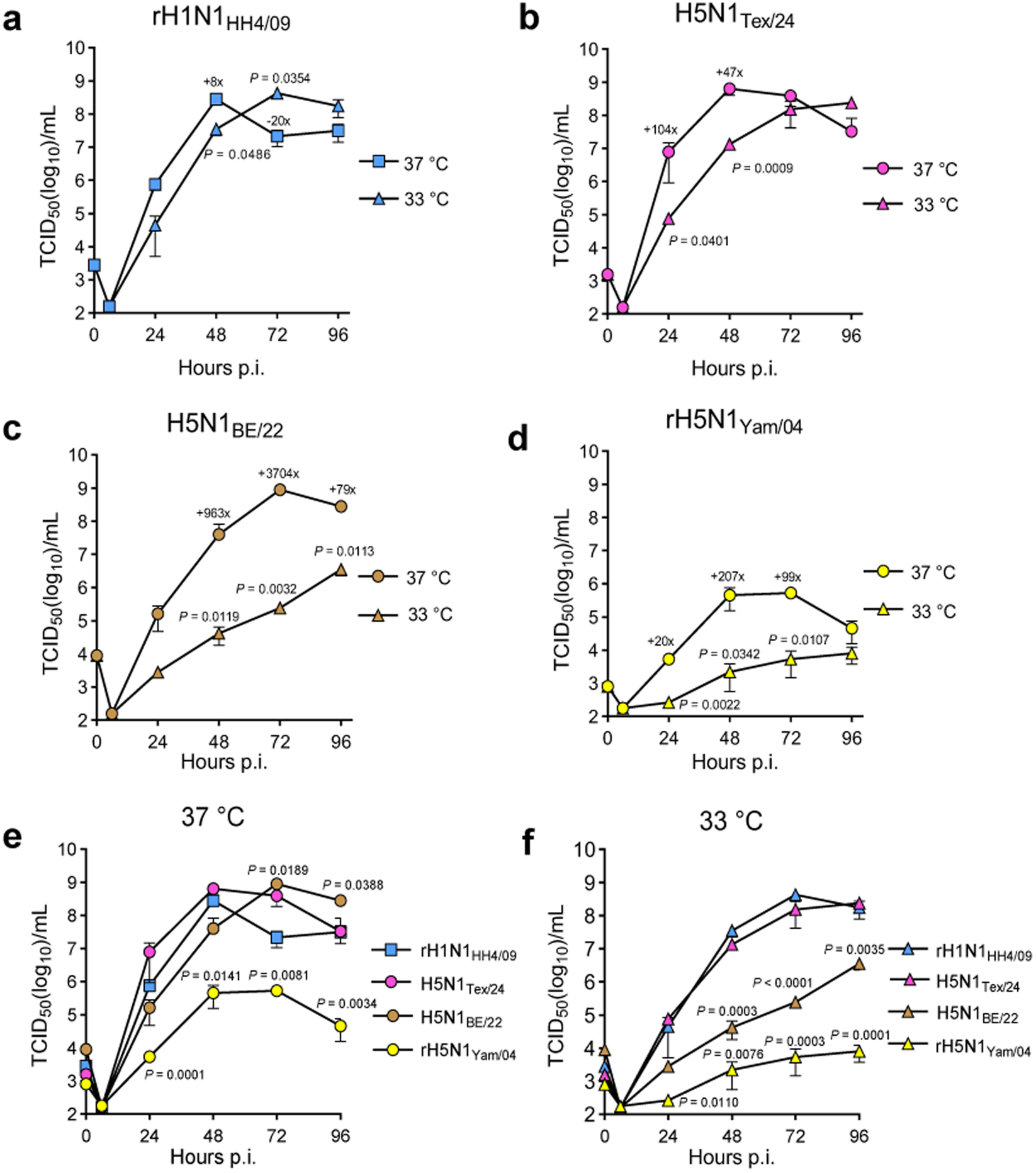
Temperature-sensitive replication of influenza A viruses in HNE cultures. **a – d** Human nasal epithelial (HNE) cells differentiated at the air-liquid interface were maintained at either 37 °C or 33 °C and infected apically with rH1N1_HH4/09_ (**a**), H5N1_Tex/24_ (**b**), H5N1_BE/22_ (**c**) or rH5N1_Yam/04_ (**d**) at a multiplicity of infection (MOI) of 0.001. At the indicated time points, apical washes were collected and infectious virus titers determined by endpoint titration on MDCK cells. **e, f** Comparison of multi-step virus replication kinetics at 37 °C (**e**) and 33 °C (**f**). Data represent mean infectious titers ± SD from three infection experiments. Statistical analyses were performed using two-way ANOVA with Tukey’s multiple comparisons test (**a – d**) or Dunnett’s multiple comparisons test using the rH1N1_HH4/09_ curve as reference (**e, f**). Only statistically significant *P* values are shown. The relative increase or decrease in infectious titers at specific time points in the 37 °C replication curves compared with the corresponding 33 °C curves is indicated (**a – d**).

### Amino acid substitutions at PB2 position 631 and PA position 497 affect H5N1_Tex/24_ replication at 33 °C but not 37 °C

Avian influenza viruses replicate most efficiently at temperatures between 41 °C and 42 °C, which is the body temperature of their avian hosts^38^. To be able to replicate efficiently in the human upper respiratory tract, avian influenza viruses must also adapt to the lower temperatures prevailing in this tissue. A characteristic mutation that frequently arises when avian influenza viruses adapt to mammals is PB2-E627K^60^. This mutation enhances interaction with cellular ANP32 proteins and thereby increases RNA polymerase activity^29^. However, it has also been reported that the same mutation enhances the replication of avian influenza viruses at lower temperatures^26,44^. Interestingly, in H5N1_Tex/24_, as in most other bovine H5N1 isolates, the enhanced interaction with ANP32 proteins is not mediated by PB2-E627K but rather by the amino acid substitutions PB2-M631L and PA-K497R^35^. Interestingly, neither PB2-E627K nor PB2-M631L and PA-K497R are present in H5N1_BE/22_, which replicates less efficiently than H5N1_Tex/24_ at 33 °C (**Fig. 2c**).

To investigate the role of the specific amino acid residues at position 631 of PB2 and position 497 of PA, respectively, we generated recombinant mutant viruses and compared their replication in HNE cultures at 37 °C and 33 °C with that of the parental viruses (**Fig. 3**). As observed before (see **Fig. 2b**), the parental rH5N1_Tex/24_ replicated efficiently at 33 °C although there was some delay at 48 h and 72 h compared to 37 °C **(Fig. 3a).** In contrast, the mutant rH5N1_Tex/24_(PB2_L631M_/PA_R497K_) replicated at 33 °C to significantly lower infectious titers than at 37 °C (**Fig. 3b**). Direct comparison of parental and mutant H5N1_Tex/24_ showed that replication of both viruses is comparable at 37 °C (**Fig. 3c**), while rH5N1_Tex/24_(L631M/R497K) replicated less efficiently than rH5N1_Tex/24_ at 33 °C, although only titers at 96 h p.i. differed significantly (**Fig. 3d**).

**Fig. 3.**
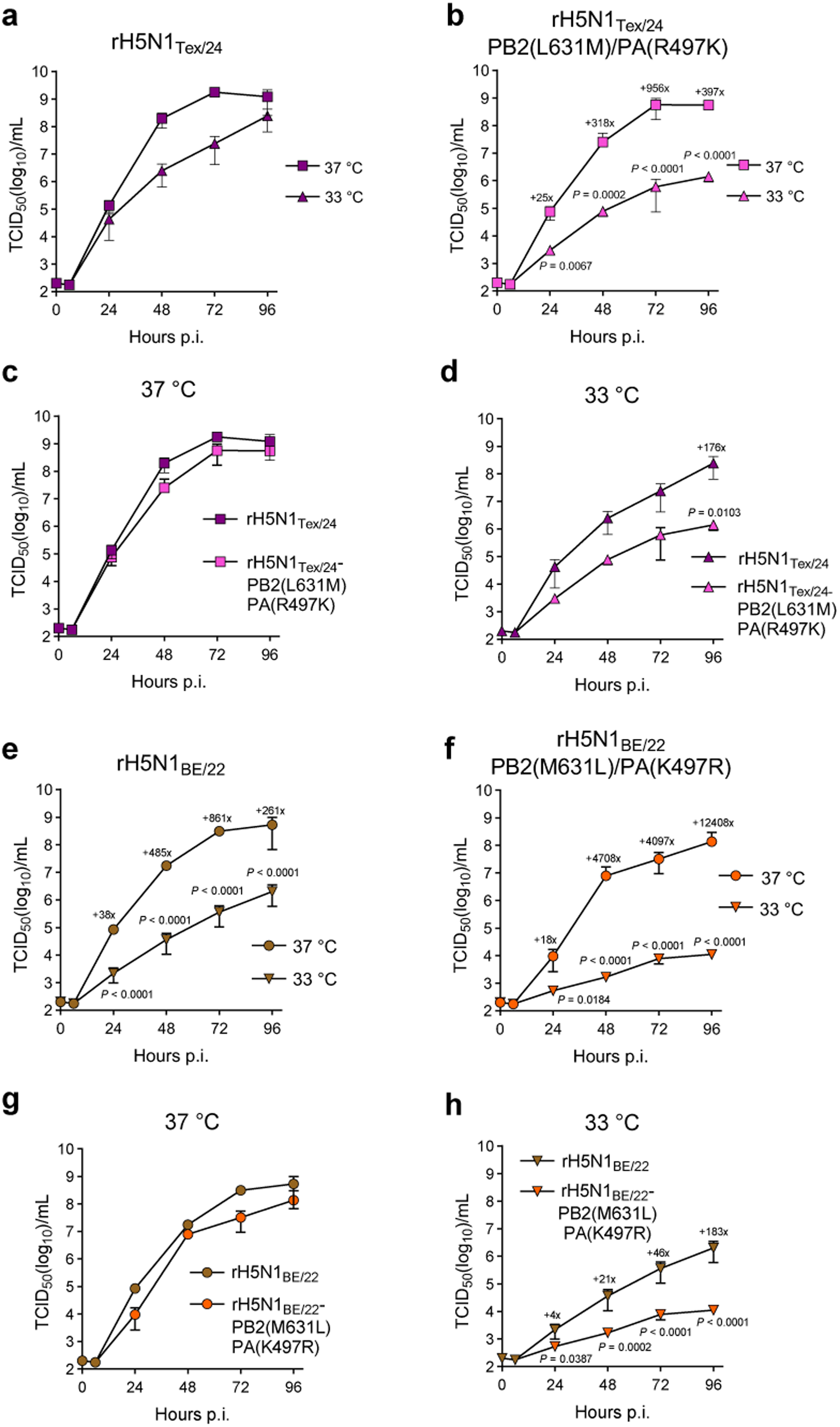
Impact of amino acid substitutions in PB2 and PA on temperature-sensitive replication of influenza A viruses. **a, b** Human nasal epithelial (HNE) cultures differentiated at the air-liquid interface were infected apically with either rH5N1_Tex/24_ (**a**) or H5N1_Tex/24_PB2(L631M)/PA(R497K) (**b**) at a multiplicity of infection (MOI) of 0.001 and maintained at either 37 °C or 33 °C for 96 h. **c, d** Comparison of multi-step replication kinetics of the indicated viruses at 37 °C (**c**) and 33 °C (**d**). **e, f** HNE cultures were infected with either H5N1_BE/22_ (**c**) or rH5N1_Yam/04_ (**d**) at an MOI of 0.001 and maintained at either 37 °C or 33 °C for 96 h. **g, h** Comparison of multi-step replication kinetics of the indicated viruses at 37 °C (**g**) and 33 °C (**h**). At the indicated time points, apical washes were collected and infectious titers determined by endpoint titration on MDCK cells. Data represent mean infectious titers ± SD from three infection experiments. Statistical analyses were performed using two-way ANOVA with Tukey’s multiple comparisons test. Exact *P* values for significant differences are indicated in the graphs. Relative increase or decrease in infectious titers at selected time points between the compared replication curves are shown.

Similar studies were conducted with rH5N1_BE/22_ and the mutant H5N1_BE/22_(M631L/K497R), which exhibited the reverse amino acid substitutions, PB2-M631L and PA-K497R. The parental virus H5N1_BE/22_ replicated to significantly lower titers at 33 °C compared to 37 °C (**Fig. 3e**), which was consistent with the temperature-dependent replication of the original H5N1-BE/22 isolate (**Fig. 2c**). Surprisingly, the introduction of the two amino acid substitutions PB2-M631L and PA-K497R did not lead to improved replication at 33 °C but rather reduced virus replication at this temperature (**Fig. 2f, h**). In contrast, PB2-M631L and PA-K497R had no effect on replication of rH5N1_BE/22_ at 37 °C (**Fig. 2g**). Collectively, these findings suggest that the substitutions PB2-M631L and PA-K497R contribute to efficient replication at lower temperatures, but only in the genetic background of H5N1_Tex/24_. In the genetic context of H5N1_BE/22_ the reverse mutations had no beneficial effect on replication at 33 °C or even worsened replication further, suggesting that adaptation to lower temperatures is most likely a polygenic and context-dependent trait.

### H5N1_Tex/24_ infection of HNE cells does not lead to IFN-λ secretion at the temperature of the upper airways

Epithelial cells can sense viruses, triggering the transcription of IFN-λ genes. These cytokines are important components of the innate immune system of epithelial cells^62^. Binding of IFN-λ to cognate receptors on epithelial cells triggers intracellular signaling pathways that lead to the transcription of numerous genes, many of which encode proteins with antiviral activity such as human myxovirus resistance protein 1 (MxA)^47^. Since H5N1_Tex/24_ can replicate efficiently in differentiated HNE cells at both 33 °C and 37 °C (see **Fig. 2**), this virus may be capable of evading the innate immune response. To investigate whether H5N1_Tex/24_ infection would lead to the induction and release of interferons, the basolateral cell culture medium derived from infected HNE ALI cell cultures was tested for IFN-λ1/3 and IFN-β by ELISA. At 33 °C, significant amounts of IFN-λ1/3 were released only at 72 h and 96 h post infection with rH1N1_HH4/09_, while infection with H5N1_Tex/24_ or H5N1_BE/22_ did not lead to significantly elevated IFN-λ levels at any time point (**Fig. 4a**). At 37 °C, increased levels of IFN-λ1/3 were detected at 48 h post infection with rH1N1_HH4/09_, while infection with H5N1_Tex/24_ or H5N1_BE/22_ led to increased levels of IFN-λ1/3 only at 72 h and 96 h post infection (**Fig. 4b**). For IFN-β, a very similar pattern of release was found at 33 °C and 37 °C (**Supplementary Fig. 2a, b**).

**Fig. 4.**
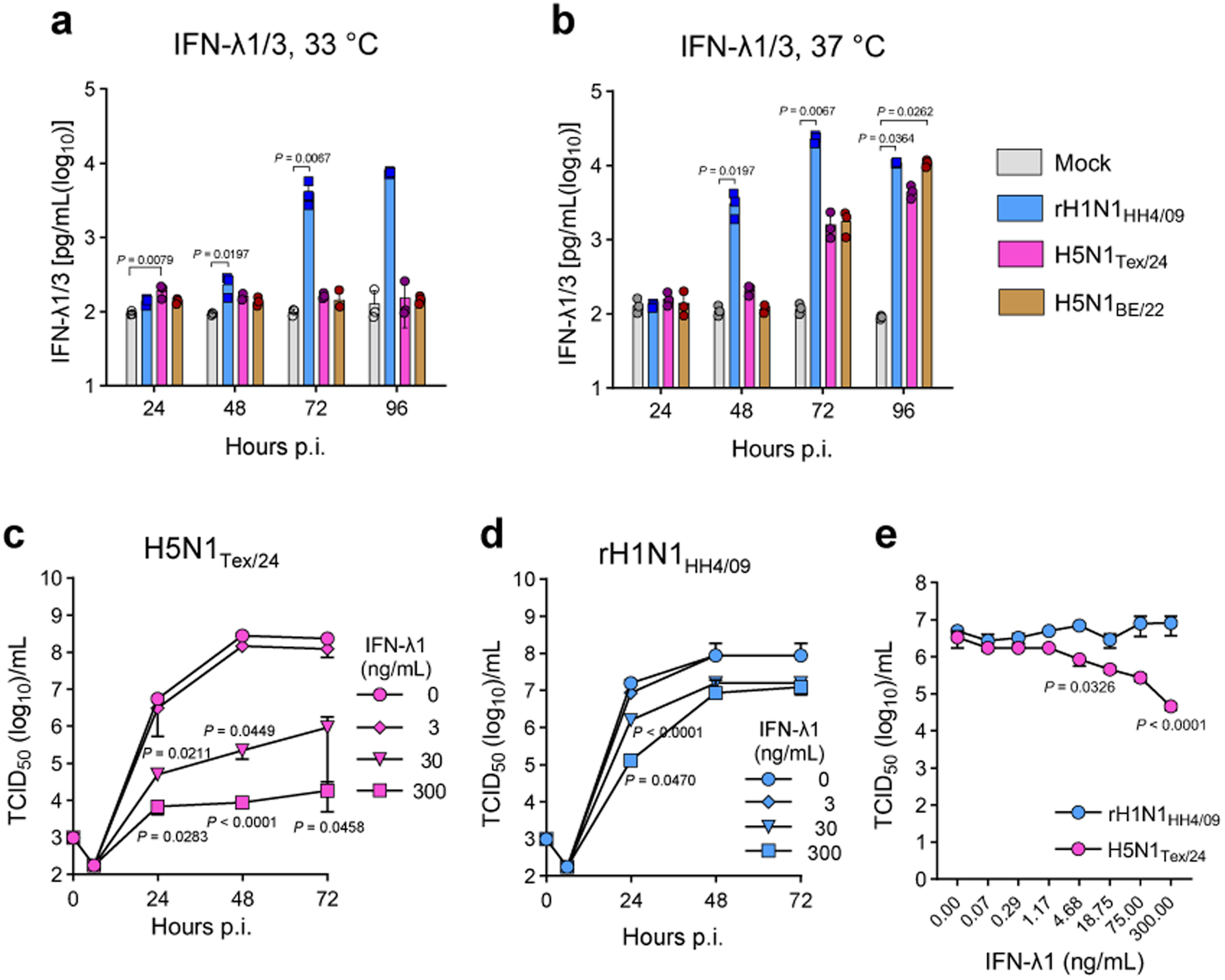
H5N1_Tex/24_ is sensitive to the antiviral activity of human IFN-λ1. **a, b** Human nasal epithelial (HNE) cultures differentiated at the air-liquid interface were infected with the indicated viruses at a multiplicity of infection (MOI) of 0.001 and maintained at either 33 °C (**a**) or 37 °C (**b**) for 96 h. At the indicated time points, basolateral medium was collected and IFN-λ1/3 levels were quantified by ELISA. **c, d** HNE air-liquid interface cultures were pretreated basolaterally for 24 h with medium containing the indicated concentrations of human IFN-λ1 and subsequently infected apically with either H5N1_Tex/24_ (**c**) or rH1N1_HH4/09_ (**d**) at an MOI of 0.001. At the indicated time points, apical washes were collected and infectious virus titers in the wash fluid determined. **e** A549 cells were treated for 24 h with the indicated of human IFN-λ1 and subsequently infected with either H5N1_Tex/24_ or rH1N1_HH4_ at an MOI of 0.01. Infectious virus titers in the cell culture supernatants were determined at 24 h post infection. Data are presented as mean titers ± SD from three infection experiments. Statistical significance was determined by non-parametric Kruskal-Wallis with Dunn′s multiple comparisons test using mock-infected cells as reference (**a, b**) or using the two-way ANOVA with Tukey′s multiple comparisons test (**c – e**). Only statistically significant *P* values are shown.

### H5N1_Tex/24_ is sensitive to the antiviral effects of IFN-λ

To assess the sensitivity of H5N1_Tex/24_ and rH1N1_HH4/09_ to IFN-λ-induced antiviral activity, well differentiated HNE ALI cultures were pretreated for 24 h at 37 °C with IFN-λ1 (3, 30, or 300 ng.mL⁻¹) added to the basolateral compartment or were left untreated. Cultures were subsequently infected from the apical side with H5N1_Tex/24_ or rH1N1_HH4/09_ at an MOI of 0.001. Apical washes were collected at 24, 48, and 72 h p.i., and infectious virus titers were determined. Pretreatment with 30 ng.mL⁻¹ IFN-λ1 significantly inhibited H5N1_Tex/24_ replication at all time points, resulting in reduction of infectious titers by 2.0 log₁₀ at 24 h, 3.1 log₁₀ at 48 h and 2.4 log₁₀ at 72 h p.i.. Increasing the IFN-λ1 concentration to 300 ng.mL⁻¹ led to an even more pronounced suppression of viral replication, with titer reductions of 2.9, 4.5 and 4.1 log₁₀ at 24, 48 and 72 h, respectively (**Fig. 4c**). By contrast, replication of rH1N1_HH4/09_ was only modestly affected by IFN-λ1 pretreatment, showing reductions of 1.0 log₁₀ and 2.0 log₁₀ at 24 h p.i. with 30 and 300 ng.mL⁻¹ IFN-λ1, respectively, whereas differences at later time points were not statistically significant (**Fig. 4d**). To corroborate these findings in an additional cellular system, A549 cells were pretreated for 24 h with IFN-λ1 (up to 300 ng.mL⁻¹) and subsequently infected with H5N1_Tex/24_ or rH1N1_HH4/09_ at an MOI of 0.01. Quantification of infectious virus released into the supernatant at 24 h p.i. revealed a dose-dependent inhibition of H5N1_Tex/24_ replication by IFN-λ1, whereas replication of rH1N1_HH4/09_ remained largely unaffected (**Fig. 4e**). Together, these results demonstrate that the bovine-derived H5N1_Tex/24_ virus is markedly more sensitive to the antiviral effects of IFN-λ1 than the pandemic rH1N1_HH4/09_ virus.

### HNE cells express both avian- and human-type influenza virus receptors

The receptor-binding specificity of influenza virus hemagglutinin (HA) is a key determinant of host range and tissue tropism^17^. Previous studies have shown that bovine-derived H5N1 viruses preferentially bind α2,3-linked sialic acids, characteristic of avian-type receptors^20^, although low affinity binding to α2,6-linked sialic acid, the predominant human-type receptor, has also been reported^63^. To define the distribution of influenza virus receptors in differentiated HNE and human bronchial epithelial (HBE) ALI cultures, we took advantage of biotin-labeled *Sambucus nigra* agglutinin (SNA), which recognizes α2,6-linked sialic acids^64^, and biotin-conjugated MAL-I/II lectins from *Maackia amurensis* which selectively bind α2,3-linked sialic acids^65^. In HNE cultures, both receptor types were abundantly expressed, with 58% of cells positive for α2,3-linked sialic acids and 72% positive for α2,6-linked sialic acids (**Supplementary Fig. 3a, b**). Similarly, HBE cultures exhibited widespread receptor expression, with α2,3-linked sialic acids detected in 71% of cells and α2,6-linked sialic acids in 96% of cells (**Supplementary Fig. 3c, d**). Given the high prevalence of both receptor types, a substantial proportion of epithelial cells is likely to co-express α2,3- and α2,6-linked sialic acids. Together, these findings indicate that receptor availability in HNE ALI cultures is unlikely to constitute a major barrier to infection by avian influenza A viruses.

Previous studies have demonstrated that during a single-cycle infection of differentiated human tracheobronchial epithelial cells, human influenza viruses preferentially infect non-ciliated cells, whereas avian viruses mainly infect ciliated cells. This pattern correlates with the predominant localization of receptors for human viruses (α2,6-linked sialic acids) on non-ciliated cells and of receptors for avian viruses (α2,3-linked sialic acids) on ciliated cells^66^. To analyse the specific tropism of human and avian influenza viruses for ciliated or non-ciliated nasal epithelial cells, differentiated HNE cultures were infected with either rH1N1_HH4/09_, H5N1_Tex/24_, H5N1_BE/22_, or rH5N1_Yam/04_ (MOI of 1) and fixed with formalin 8 h p.i. Immunofluorescence analysis of the cells by confocal laser scanning microscopy revealed that a majority of the HNE cells are ciliated cells and express both α2,3- and α2,6-linked sialic acids (**Fig. 5**), suggesting that ciliated nasal epithelial cells are targeted by both human and avian influenza viruses.

**Fig. 5.**
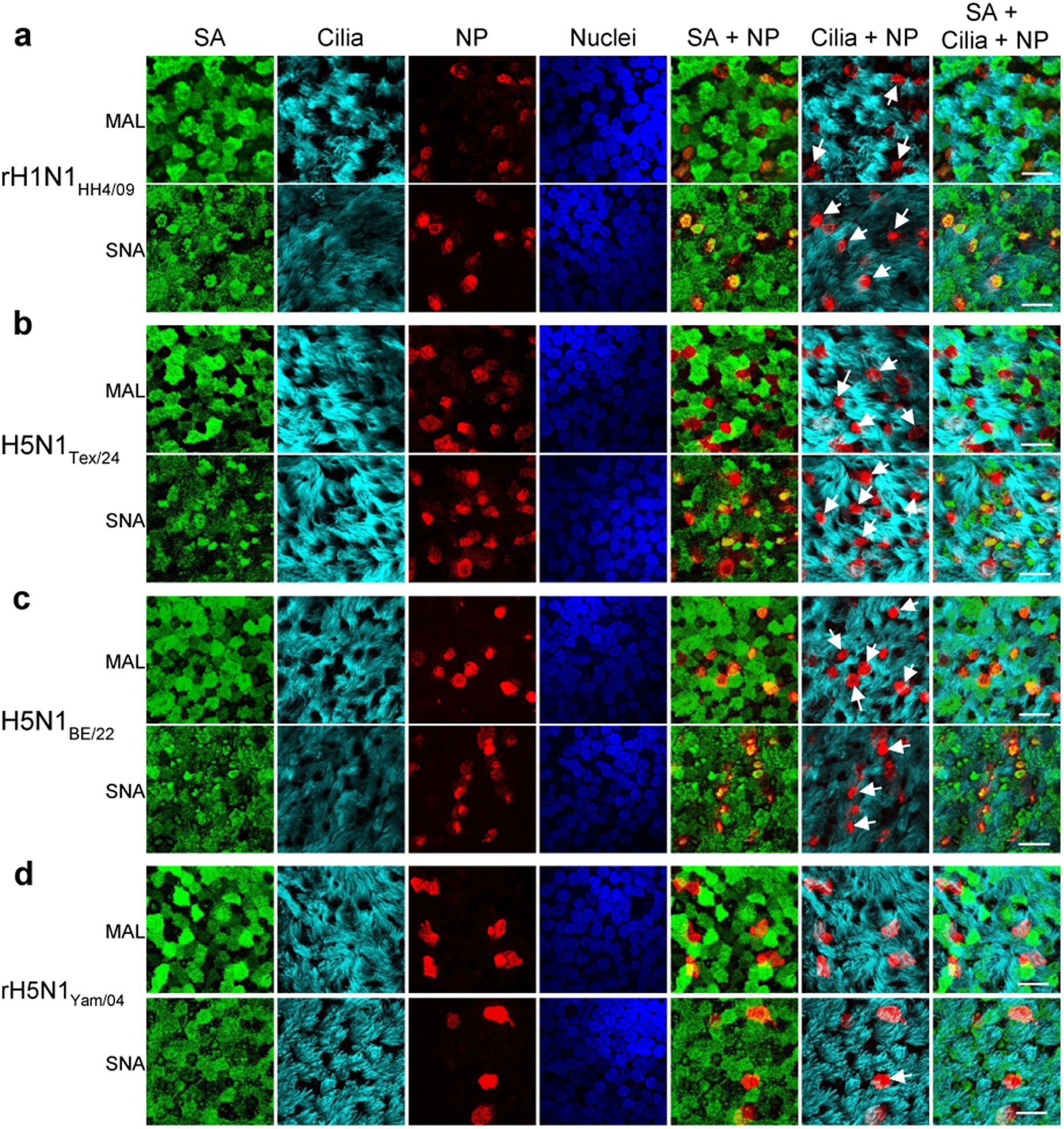
Detection of avian- and human-type receptors in HNE cells. **a - d** Human nasal epithelial (HNE) cultures differentiated at the air-liquid interface were infected with either rH1N1_HH4/09_ (**a**), H5N1_Tex/24_ (**b**), H5N1_BE/22_ (**c**) or rH5N1_Yam/04_ (**d**) at an MOI of 1. Cells were incubated at 37 °C for 8 h and subsequently fixed with formalin. Non-permeabilized cells were stained with *Maackia amurensis* lectin (MAL) to detect α2,3-linked sialic acid (SA) or with *Sambucus nigra* agglutinin (SNA) to detect α2,6-linked SA. Ciliated cells were identified using an antibody against acetylated tubulin. Influenza virus-infected cells were detected following Triton X-100 permeabilization using a monoclonal antibody against the conserved nucleoprotein (NP) antigen. Nuclei were stained with DAPI. Images were acquired by confocal laser scanning microscopy following indirect immunofluorescence staining. Scale bar, 25 µm. Representative images from two infection experiments are shown. White arrows point to infected cells lacking cilia.

### H5N1_Tex/24_ replication in HNE cultures is associated with rapid cilia loss

Ciliated epithelial cells are critical for airway defense by mediating mucociliary clearance of mucus and inhaled particles. Immunofluorescence analysis revealed that the majority of HNE cells infected with rH1N1_HH/09_, H5N1_Tex/24_ or H5N1_BE/22_ displayed either complete loss or a marked reduction of cilia, as indicated by white arrows in **Fig. 5**. To further assess the effect of H5N1_Tex/24_ infection on ciliated cells, differentiated HNE cultures were infected with H5N1_Tex/24_ or rH1N1_HH4/09_ at an MOI of 0.1. Mock-infected HNE cultures contained a high proportion of ciliated epithelial cells (> 82%) (**Fig. 6a, b**). At 16 h p.i. with rH1N1_HH4/09_, the proportion of ciliated epithelial cells remained largely unchanged, whereas at 24 h p.i. it decreased to 64%, corresponding to an 18% reduction (**Fig. 6b**). In contrast, following H5N1_Tex/24_ infection, the proportion of ciliated epithelial cells had already declined to 64% at 16 h p.i. and further decreased to 41% at 24 h p.i., representing an overall reduction of approximately 50%. These findings indicate that cilia loss progressed substantially more rapidly in H5N1_Tex/24_-infected cultures than in rH1N1_HH4/09_-infected cultures. Infected cells also showed enhanced caspase 3 activity consistent with apoptosis, which became more pronounced between 16 h to 24 h p.i., particularly in H5N1_Tex/24_-infected HNE cultures (**Fig. 5a**). Taken together, these results show that infection with H5N1_Tex/24_ infection is associated with extensive deciliation of cells, suggesting infection with this virus has the potential to severely impair mucociliary clearance.

**Fig. 6.**
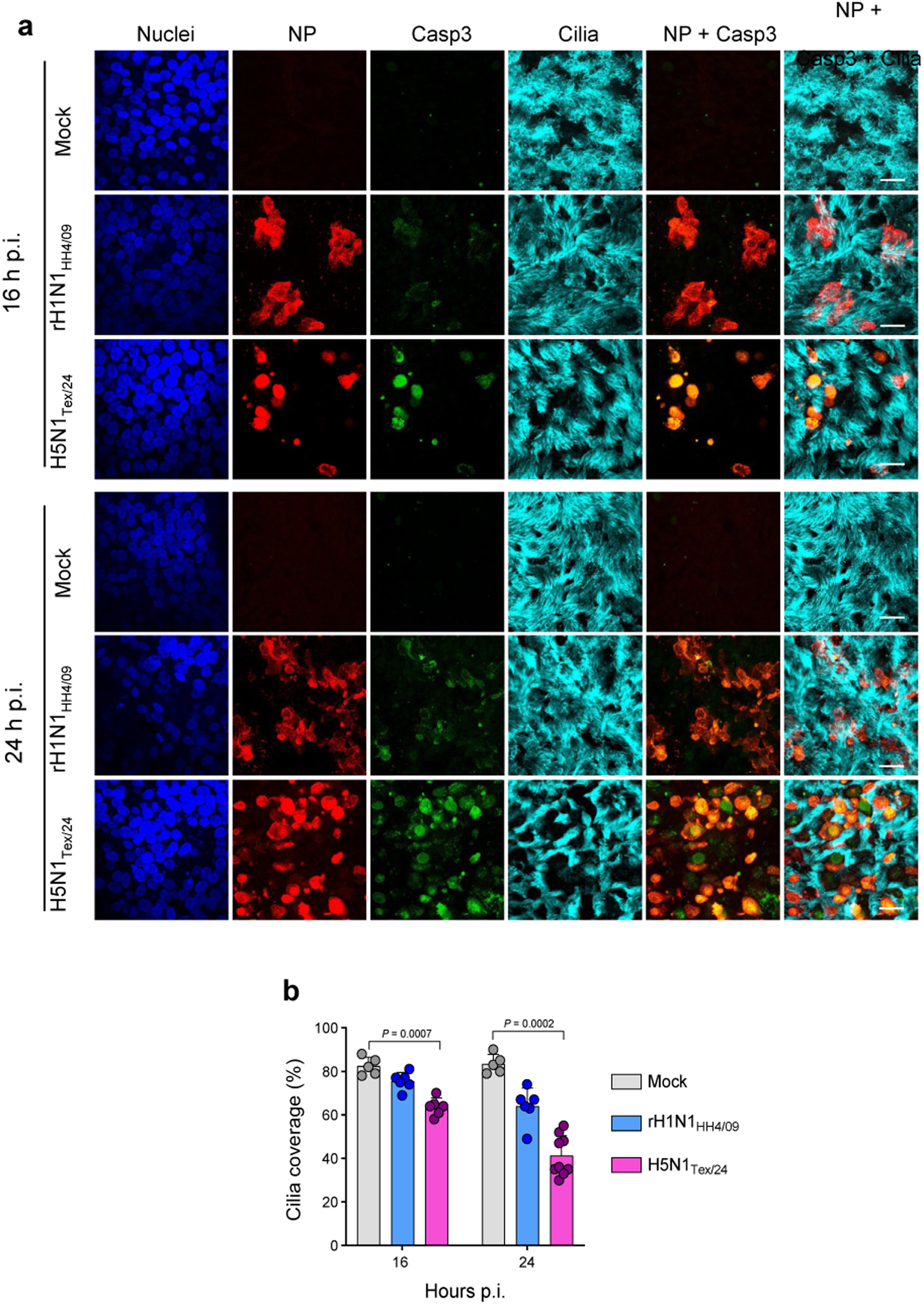
Infection of HNE cultures induces apoptosis and loss of cilia. **a** HNE cells differentiated at the air-liquid interface were mock-infected or infected with H5N1_Tex/24_ or rH1N1_HH4/09_ using an MOI of 0.1. At 16 h and 24 h p.i., cells were incubated with a fluorogenic caspase-3 substrate and subsequently fixed with formalin. Influenza virus-infected cells were stained for NP antigen, cilia were labelled with an antibody against acetylated tubulin, and nuclei were stained with DAPI. Scale bar, 25 µm. **b** Quantification of cilia coverage at 16 h p.i. and 24 h p.i. using ImageJ. Five to nine images per condition, each containing approximately 100 cells, were analyzed. Statistical significance was determined by Kruskal-Walllis with Dunn′s multiple comparisons test. Only statistically significant *P* values are shown.

## Discussion

H5N1 HPAI viruses of clade 2.3.4.4b have spread globally in recent years, causing mass mortality in wild birds and significant losses in domestic poultry^67^. These viruses have also crossed species barriers and infected a growing number of mammalian species, including humans^3^. Most human cases have occurred after close contact with infected poultry or, more recently, infected dairy cows^6,9,12,68^. Reported human disease has been mild in most instances^9^, and human-to-human transmission has not been documented to date^3,69^. One proposed explanation is that these avian-origin viruses cannot replicate efficiently in the human upper respiratory tract, owing to factors such as limited availability of compatible receptors, suboptimal temperature for replication, or insufficient adaptation to evade innate immune responses^16^. Using primary, well differentiated HNE cells cultured at the air–liquid interface, we show here that a bovine-derived H5N1 virus replicates as efficiently as the 2009 pandemic H1N1 virus in the human nasal epithelium. Interestingly, H5N1_Tex/24_ replicated more efficiently in HNE cells when an MOI of 0.001 was used for infection. In contrast, infection with higher viral doses (MOI of 0.01 or 0.1) resulted in flatter growth curves (see **Fig. 1**), similar to recently published data^24^. Surprisingly, this phenomenon was not observed when recombinant viruses were used, leading us to assume that this effect stems from defective interfering particles, which were present in the viral isolate but not in the recombinant viruses^70^.

Adaptation of avian influenza viruses to mammals often involves changes in the viral RNA polymerase complex^60,71^. The PB2-E627K substitution, a canonical marker of mammalian adaptation, enhances polymerase activity via improved interaction with ANP32 proteins^72-74^. However, only a minority of human-derived clade-2.3.4.4b viruses sequenced to date carry PB2-E627K^33,75^. H5N1_Tex/24_ like most other bovine H5N1 isolates lack PB2-E627K as well as other mammalian-adaptive PB2 substitutions such as PB2-Q591R^29^, PB2-D701N^28,76^, and PB2-T271A^77^ (**Table 2**). The majority of bovine H5N1 viruses including H5N1_Tex/24_ contain the adaptive mutation PB2-M631L which has been shown to improve polymerase activity in human cells^12,33^. PB2-M631L maps to the polymerase-ANP32 interface and enhances viral RNA polymerase activity by supporting interaction with bovine ANP32 proteins^35^. Polymerase activity is further enhanced by the adaptive PA-K497R substitution, which is present in most bovine-derived H5N1 viruses^35^. Interestingly, rH5N1_Tex/24_ carrying the reverse mutations PB2-L631M and PA-R497K replicated as efficiently as the parental virus in HNE cultures at 37 °C. Moreover, rH5N1_BE/22_, which lacks these adaptive mutations, and the corresponding mutant rH5N1_BE/22_(PB2-L631M/PA-K497R) replicated well on HNE cells at 37 °C, indicating that the PB2-M631L and PA-K497R mutations might be of minor importance for replication on HNE cells. In this context, it is noteworthy to mention that the PB2-M631L and PA-K497R mutations are also absent from bovine H5N1 viruses that belong to genotype D1.1, suggesting that other, so far unknown adaptive mutations may exist in the polymerase genes of these viruses that support replication in mammalian hosts.

One mutation that did arise during replication in HNE cells was PA-I38M (**Table 1**), which confers resistance to baloxavir^78,79^. Although present in only ∼5% of reads, its rapid emergence and lack of an apparent fitness cost in human cells is concerning, underscoring that clade 2.3.4.4b viruses can readily acquire antiviral-resistance mutations.

Temperature represents an important barrier to mammalian adaptation^39,42^. Avian influenza viruses typically replicate optimally at 41-42 °C, the body temperature of birds^38^, but typically show restricted replication at the lower temperatures of the human upper airways. Here, bovine H5N1_Tex/24_ replicated efficiently at both 33 °C and 37 °C, whereas H5N1_BE/22_, despite belonging to the same clade, replicated poorly at 33 °C. The PB2-E627K substitution, which is known to promote replication at lower temperatures^44,80^, is absent in H5N1_Tex/24_ (**Table 2**). This suggests that other genetic determinants are responsible for efficient virus replication at 33 °C. Our experiments with recombinant H5N1_Tex/24_ mutants demonstrate that the reverse mutations PB2-L631M and/or PA-R497K do not affect replication of H5N1_Tex/24_ at 37 °C but have an attenuating effect at 33 °C. These results imply that these amino acid positions 631 in PB2 and 497 in PA are critical for virus replication at suboptimal temperature by increasing polymerase activity beyond a critical threshold. Surprisingly, the avian H5N1_BE/22_ isolate did not show enhanced replication at 33 °C when the PB2-M631L and PA-K497R substitutions were introduced, suggesting that the genetic background is also critical in this regard. Identifying the determinants that enhance the replication of bovine-derived H5N1 virus at 33 °C will be an important direction for future work.

The receptor specificity of the HA protein is considered a key determinant of the host tropism of influenza viruses; furthermore, the availability of suitable receptors in the respiratory tract acts as a major species barrier to infection of new hosts^81^. While 2009 pandemic H1N1 viruses bind predominantly to α2,6-linked sialic acids^56,57^, recent studies indicate that bovine H5N1 viruses retain a largely avian receptor-binding profile and preferentially recognize α2,3-linked sialic acids^20,21^. Although a single mutation at HA position 222 (H5 numbering) can shift binding toward α2,6-linked sialic acids^22^, we did not detect this mutation in the H5N1_Tex/24_ isolate (**Table 2**), neither before nor after replication for 72 h on HNE cells (**Table 1**). Lectins staining demonstrated that differentiated HNE cells display both α2,3- and α2,6-linked sialic acids, consistent with the previous finding that the human respiratory tract has potential binding sites for human and avian influenza viruses^82^. These findings also align with a recent immunohistochemical study showing efficient HA binding of bovine H5N1 viruses to human conjunctival, tracheal, and mammary tissues^63^. A recent study comparing receptor-binding of contemporary clade 2.3.4.4b with 2005 clade 2.1.3.2 H5N1 viruses suggests that five amino acid substitutions located near the HA receptor-binding site of clade 2.3.4.4b H5N1 viruses may increase receptor affinity but not specificity^24^. Together, these observations indicate that receptor distribution in differentiated primary HNE cells does not impose a strong barrier to infection and thus provides limited selective pressure for adaptation toward α2,6-linked receptors.

Innate immune restriction represents an additional barrier to mammalian infection by animal influenza viruses^73^. Our data suggest that - in contrast to rH1N1_HH4/09_ - H5N1_Tex/24_ and H5N1_BE/22_ induce very little IFN-β and IFN-λ1/3 in differentiated HNE cultures, in line with a previous report^24^. Remarkably, the H5N1 viruses did not induce any IFN in HNE cells that were grown at 33 °C, while they did induce IFN when the cells were grown at 37 °C, but only at late time points (see **Fig. 4a, b**). These findings suggest that both H5N1_Tex/24_ and H5N1_BE/22_, but not rH1N1_HH4/09_, may actively suppress the synthesis of IFNs. Indeed, there is evidence that clade 2.3.4.4b viruses have a pronounced host shut-off activity^31^.

The most important viral factors known to contribute to the prevention of IFN induction are the NS1 protein^83,84^ and the PA-X protein^85-87^, both proteins acting in a concerted manner^88-90^. All three H5N1 viruses analysed in this study harbor the mutations NS1-L103F and NS1-I106M (**Table 2**), which have been linked to enhanced binding of NS1 to the pre-mRNA processing factor CPSF30^91^. All three H5N1 viruses also harbor the PA-X-T118I and PA-X-I127V substitutions which are associated with increased PA-X-mediated host shut-off activity^92^. There are many additional amino acid differences between the PA-X proteins of the clade 2.3.4.4b viruses H5N1_Tex/24_ and H5N1_BE/22_ compared with H5N1_Yam/04_, but it is not yet known whether any of these differences affect the shut-off activity of the viruses in human cells. Future studies must show whether the low replication efficacy of H5N1_Yam/04_ is due to lower host shut-off activity and increased induction of IFNs.

Influenza viruses can evade the antiviral activity of interferon-stimulated genes (ISG). For example, certain mutations in the nucleoprotein (NP) can overcome the restriction imposed by MxA and BTN3A3^54,93^. It is noteworthy that the NP proteins of H5N1_Tex/24_ and H5N1_BE/22_ but not rH5N1_Yam/04_ have the NP-Y52H substitution (**Table 2**), which confers resistance to the restriction factor BTN3A3^94^. However, H5N1_Tex/24_ lacks known MxA resistance mutations such as NP-F313Y and NP-N319K^94,95^, and remains susceptible to the antiviral activity of human MxA^47^. H5N1_Tex/24_ induces only low levels of type I and type III IFNs in HNE cells, perhaps allowing the virus to tolerate some sensitivity to ISGs. In contrast, rH1N1_HH4/09_ has characteristic mutations in the NP protein that confer resistance to MxA^54^.

Using an antibody against acetylated tubulin, we detected in differentiated HNE cultures a high proportion of ciliated epithelial cells (see **Fig. 5**). The cilia of these cells represent hair-like organelles that project from the apical cell surface and basically consist of a centriole-derived, microtubule core, the axoneme^96^. In our experiments, infected HNE cells seem to have no or little cilia already at 8 h p.i. (**Fig. 5**). Although it cannot be excluded that both rH1N1_HH4/09_ and H5N1_Tex/24_ were specifically targeting non-ciliated epithelial cells, it is more likely that infection caused the loss of cilia because of the virus-mediated host shut-off and the induction of apoptosis^97^. There was a more pronounced loss of cilia linked to infection with H5N1_Tex/24_ compared to infection with rH1N1_HH4/09_ which might be due to a faster progressing apoptosis in H5N1_Tex/24_-infected cells. The loss of cilia can have a deleterious effect on mucociliary clearance and disease severity^98^.

Overall, our results show that bovine-derived H5N1_Tex/24_ replicates at high titers in primary human nasal epithelial cells even though it lacks many canonical markers of adaptation to mammals. This observation contrasts with the currently low number of confirmed human infections. There are several possible explanations for this discrepancy. First, the sensitivity of H5N1_Tex/24_ to the antiviral effects of MxA may limit virus dissemination to the lower respiratory tract and thus disease severity. Second, human infections may go undetected because they are completely asymptomatic or because they are associated with only mild symptoms. Third, efficient airborne transmission of H5N1 requires that HA induces membrane fusion at pH values significantly lower than pH 6.0^99,100^. However, the HA protein of bovine H5N1 retains typical avian characteristics, with fusion triggered at approximately pH 6.0^21,101^. Finally, pre-existing immunity to the 2009 pandemic H1N1 virus have been shown to provide at least partial protection of ferrets from infection by bovine H5N1 virus^102-105^. There is also pre-existing cross-reactive immunity to highly pathogenic avian influenza 2.3.4.4b A(H5N1) virus in the human population^105,106^. In particular, cross-reactive antibodies against the NA protein of human H1N1 viruses can particularly inhibit avian N1 sialidase activity^107^, thereby potentially limiting replication of clade-2.3.4.4b viruses. The recent fatal human case caused by a H5N5 HPAI virus in the United States is alarming^108^, as there is probably no pre-existing immunity to the N5 antigen in the human population. Overall, our results show that H5N1_Tex/24_ has a remarkable ability to replicate in primary human nasal epithelial cells.

## Methods

### Cells

Primary human nasal epithelial (HNE) and human bronchial epithelial (HBE) cells were obtained from Epithelix Sàrl as MucilAir™-Pool nasal and MucilAir™-Pool bronchial models, derived from pools of 14 and 5 healthy donors, respectively. Cultures displayed robust epithelial barrier integrity, with transepithelial electrical resistance (TEER) values >200 Ω·cm². Cells were confirmed negative for mycoplasma, HIV-1, HIV-2, hepatitis B virus, and hepatitis C virus. Airway epithelial tissues were maintained on porous Transwell inserts (6.5 mm diameter, 1 µm pore size) in MucilAir™ culture medium (Epithelix Sàrl), according to the manufacturer’s instructions. Each insert contained approximately 5 × 10⁵ cells.

Madin-Darby canine kidney (MDCK) type II cells were provided by Georg Herrler (University of Veterinary Medicine, Hannover, Germany) and were maintained with minimum essential medium (MEM, Thermo Fisher Scientific, Basel, Switzerland; cat. no. 31095-029) supplemented with 5% fetal bovine serum (FBS; Pan Biotech, Aidenbach, Germany; cat. no. P30-3033). Human embryonic kidney (HEK) 293T cells (American Type Culture Collection (ATCC), Manassas, USA; cat. no. CRL-3216) were maintained with Dulbecco’s Modified Eagle Medium (DMEM, Thermo Fisher Scientific; cat. no. 32430-027) supplemented with 10% FBS. A549 human lung carcinoma cells (ECACC, cat. no. 86012804), were grown in Ham’s F12 nutrient mixture (Merck KGaA, Darmstadt, Germany; cat. no. N6658) supplemented with 10% FBS. All cell lines were routinely monitored for contamination by mycoplasma.

### Viruses

A/cattle/Texas/063224-24-1/2024 (H5N1_Tex/24_) was kindly provided by Diego Diel (Cornell University, Ithaca, NY, USA). H5N1_Tex/24_ was isolated from the milk of infected dairy cows in March 2024 in Texas, USA. The virus belongs to clade 2.3.4.4b, genotype B3.13 (GISAID accession no.: EPI_ISL_19155861)^52^. H5N1_Tex/24_ virus stock was produced by two passages in MDCK cells. The complete genome sequence of this virus stock was determined and deposited in the GenBank database (accession nos.: PX831491-PX831498). A/Dalmatian pelican/Bern/1/2022 (H5N1_BE/22_) was isolated in February 2022 from a Dalmatian pelican in Bern, Switzerland. H5N1_BE/22_ virus stock was produced by passaging the virus three times in MDCK cells. The complete genomic sequence of H5N1_BE/22_ was deposited at GenBank (accession nos.: PX700809 - PX700813, PX149233 - PX149235).

Recombinant influenza viruses A/Hamburg/4/2009 (rH1N1_HH4/09_), A/chicken/Yamaguchi/7/2004 (rH5N1_Yam/04_), and A/bovine/Texas/24-029328-02/2024 (rH5N1_Tex/24_) and A/Dalmatian Pelican/Bern1/2022 (rH5N1_BE/22_) were generated using the eight-plasmid reverse genetics system^58^. cDNAs encoding the 8 genomic segments of H1N1_HH4/09_ (GenBank accession nos.: GQ166207, GQ166209, GQ166211, GQ166213, GQ166215, GQ166217, GQ166219, GQ166221) were provided by Martin Schwemmle (University of Freiburg, Germany). cDNAs encoding the eight genomic segments of H5N1_Yam/04_ (GenBank accession nos.: GU186708 - GU186715) were obtained from Yoshihiro Sakoda (Hokkaido University, Sapporo, Japan). cDNAs corresponding to the eight genomic segments of A/bovine/Texas/24-029328-02/2024 (H5N1) (GenBank accession nos.: PP599470 - PP599477) were synthesized by Genscript Biotech (Piscataway, New Jersey, USA) and cloned into the pHW2000 plasmid^58^ using the In-Fusion cloning system (Takara Bio, Saint-Germain-en-Laye, France, cat. no. 638948). Site-directed mutagenesis was used to introduce the PB2-L631M and PA-R497K substitutions into pHW2000 plasmids containing segments 1 and 3 of rH5N1_Tex/24_, respectively. Conversely, cDNAs encoding segments 1 and 3 of rH5N1_BE/22_ were modified to encode the reverse substitutions PB2-M631L and PA-K497R, respectively. The nucleotide sequence of each cloned segment was verified by Sanger sequencing.

### Virus replication kinetics in differentiated human airway ALI cultures

Well differentiated HNE or HBE cells grown at ALI were washed three times with Hank’s Balanced Salt Solution (HBSS; Gibco) to remove mucus prior to infection. The cells were then inoculated apically for 2 h at either 33 °C or 37 °C with 200 µL of the respective influenza A virus. Thereafter, the inoculum was removed, and the apical surface washed three times with HBSS to remove unbound viruses. The third wash was collected and kept at -70 °C (2h-sample). For subsequent time points, apical wash samples were collected by incubating the apical domain of the cells for 10 min with 200 µL of HBSS and stored at -70 °C until virus titration.

Infectious virus titers were determined on MDCK cell monolayers in 96-well culture plates by limiting dilution. To this end, cells were incubated in quadruplicates with serially diluted virus in MEM without FBS (100 μL/well). rH1N1_HH4/09_ containing a monobasic HA cleavage site was incubated in the presence of 1 µg/mL of acetylated trypsin (Merck KGaA, Darmstadt, Germany, cat. no. 4370285). After three days, the cells were fixed for 30 min at room temperature with 4% buffered formalin containing 0.1% (w/v) crystal violet. The plates were washed with tap water and dried. Virus titer was calculated using the Spearman-Kärber method and expressed as tissue culture infectious dose 50% per mL (TCID_50_/mL)^109^.

### Determination of influenza virus genome sequences

Total RNA was extracted from apical wash samples collected at 72 h p.i. using the QIAamp Viral RNA Mini Kit (Qiagen, Hilden, Germany; cat. no. 52904), according to the manufacturer’s instructions. RNA concentration was determined using a Qubit 4.0 Fluorometer with the Qubit RNA Broad Range and High Sensitivity Assay Kits (Thermo Fisher Scientific, cat. nos. Q10211 and Q32855), and RNA integrity was assessed on an Advanced Analytical Fragment Analyzer System using the Fragment Analyzer RNA Kit (Agilent, cat. no. DNF-471).

The eight influenza A virus genomic segments were reverse-transcribed and amplified using the SuperScript IV One-Step RT-PCR System (Thermo Fisher Scientific; cat. no. 12594100) with the Opti primer set^110^. Resulting amplicons were purified and quantified using the Qubit dsDNA High Sensitivity Assay Kit (Thermo Fisher Scientific; cat. no. Q32854) on a Qubit 4.0 Fluorometer. Amplicon size distributions were verified using an Agilent FEMTO Pulse System with the Genomic DNA 165 kb Kit (Agilent, FP-1002-0275).

Each sample PCR reaction was used as input to generate barcoded SMRTbell libraries according to the following Procedure & checklist: Preparing multiplexed amplicon libraries using SMRTbell prep kit 3.0 Procedure & checklist (PacBio, 102-359-000 REV04. Instructions in SMRT Link Sample Setup were followed to prepare the SMRTbell libraries for sequencing (PacBio SMRT Link v25) using the Revio SPRQ polymerase kit (PacBio, cat no.103-496-900) and clean-up beads (PacBio, cat. no. PN 102-158-300). The libraries were loaded at an on-plate concentration of 225 pM using adaptive loading with a 24 h movie time. All steps post RNA extraction were performed at the Next Generation Sequencing Platform, University of Bern.

### Immunofluorescence

Cells were fixed with 4% formalin for 30 min at room temperature and washed three times with PBS containing 0.1 M glycine. Following permeabilization with 0.25% (v/v) Triton X-100 (15 min, room temperature), the cells were incubated overnight at 4 °C with a monoclonal antibody directed to influenza NP (1:50 in PBS; American Type Culture Collection, Manassas, Virginia, USA, cat. no. ATCC HB-65) along with either Alexa Fluor-594-labeled mouse monoclonal antibody directed against ZO-1 (1:500; Life Technologies, Zug, Switzerland, cat. no. 339194) or with a rabbit monoclonal antibody directed against acetyl-α-tubulin (1:300; Cell Signaling, Danvers, USA, cat. no. 5335S). The unlabeled primary antibodies were detected by incubation of the cells for 1 h with goat anti-mouse IgG conjugated with Alexa Fluor-488 (1:500; Life Technologies, cat no. A11001) or anti-rabbit IgG conjugated to Alexa Fluor-647 (1:500, Life Technologies, cat. no. A21244). For detection of glycoconjugates containing α2,3-linked or α2,6-linked sialic acids, formalin-fixed, non-permeabilized cells were incubated overnight at 4 °C with biotinylated *Sambucus nigra* agglutinin (SNA, 1:200; Vector Labs, Newark, CA, USA, cat. no. B-1305-2) or combined biotinylated *Maackia amurensis* lectins I and II (Mal-I, Mal-II, 1:200 each; Vector Labs, cat nos. B-1315-2 and 1265-1). Thereafter, the cells were washed three times with PBS and incubated for 1 h with streptavidin conjugated with Alexa Fluor-488 (1:250, Life Technologies, cat. no. S11223). For staining of nuclei, cells were incubated for 5 min at room temperature with 0.1 µg/mL of 4′,6-diamidino-2-phenylindole (DAPI, Merck KGaA, Darmstadt, Germany) and mounted in ProLong Gold Antifade Mountant (Thermo Fisher). Cells were imaged using a confocal laser scanning system mounted on an ECLIPSE Ti inverted microscope (Nikon Instruments) and data processed with ImageJ software.

### Apoptosis assay

HNE cells grown in the air-liquid interface were mock-infected or infected with H5N1_Tex/24_ or rH1N1_HH4/09_ using an MOI of 0.1. At 16 h and 24 h p.i., the cells were first incubated for 1 h at 37 °C with the NucView-488 fluorogenic caspase-3 substrate (BioLegend; cat. no. 421928) at 5 µM final concentration. Subsequently, cells were fixed with 4% formalin for 30 min at room temperature. Influenza virus-infected cells were identified by immunostaining with an antibody directed against the viral NP antigen, and cilia were labelled with an antibody directed to acetylated tubulin (see above). Nuclei were counterstained with DAPI.

### Production of recombinant human IFN-λ1

The expression plasmid pcDNA3.1-IFNL1 encoding human IFNλ1 (GenBank accession no. NM_172140.2) was obtained from GenScript Biotech (Rijswijk, The Netherlands). HEK 293T cells grown in 10-cm cell culture dishes (approximately 10^7^ cells/dish) were transfected with 10 μg of plasmid DNA and 20 μl of Lipofectamin 2000 (Life Technologies, cat. no. 11668019), according to the manufacturer’s instructions. At 20 h post transfection, the cells were washed once with 10 mL of PBS and further incubated in FBS-free DMEM medium for 24 h at 37 °C. The amount of IFNλ1 in the conditioned cell culture medium was determined by ELISA (see below).

### ELISA

Levels of IFN-β and IFN-λ1/3 proteins released into the basolateral medium of HNE cells were measured with the Human IFN-β DuoSet ELISA (R&D Systems, Minneapolis, MN, USA, cat. #DY814-05) and the human IL-29/IL-28B (IFNλ1/3) DuoSet ELISA (R&D Systems, cat. no. DY1598B) according to the manufacturer’s instructions.

### Biosafety

Work with H5N1 highly pathogenic avian influenza viruses has been approved by the Swiss Federal Office of Public Health (license no. A230074-01) and was performed at IVI Mittelhäusern in laboratories complying with biosafety level 3 standards.

### Statistics

Statistical analysis was performed with GraphPad Prism 10, version 10.4.1. Unless otherwise noted, results are expressed as arithmetic means and standard deviations (SD). Specific statistical tests such as the two-way ANOVA test were employed to assess significant differences in the multi-step replication of different viruses and are indicated in the figure legends. *P* values < 0.05 were considered significant.

## Data availability

All data generated or analyzed during this study are included in this article and its supplementary information files. Source data will be accessible at Zenodo at the time of publication. The nucleotide sequences of all eight genomic segments of A/Dalmatian Pelican/Bern/1/2022 (H5N1) are available at GenBank (accession nos.: PX149233 - PX149235 and PX700809 – PX700813). Genomic sequences of A/cattle/Texas/063224-24-1/2024 (H5N1) that was propagated on MDCK cells are available at GenBank (accession nos.: PX831491 - PX831498). The PacBio sequencing reads are available in the ENA archive under project accession PRJEP106468.

## Acknowledgements

We are grateful to the Swiss Federal Food Safety and Veterinary Office (FSVO) for financial support. We thank the Next Generation Sequencing Platform of the University of Bern for performing high-throughput sequencing experiments, Martin Schwemmle and Yoshihiro Sakoda for providing plasmids and Diego Diel for providing the bovine H5N1 virus.

## Author contributions

Conceptualization: E.A.M., M.P.A., and G.Z. Investigation and data curation: E.A.M., L.B., T.D., M.W., P.G. and G.Z. Funding acquisition: M.P.A. and G.Z. Resources: C.C., S.C., C.B., V.T., M.P.A. and G.Z. Writing – original draft: E.A.M. and G.Z. Writing – review and editing: E.A.M., C.C., S.C., M.W., P.G., C.B., V.T., M.P.A., and G.Z.

---

**Supplementary Fig. 1.**
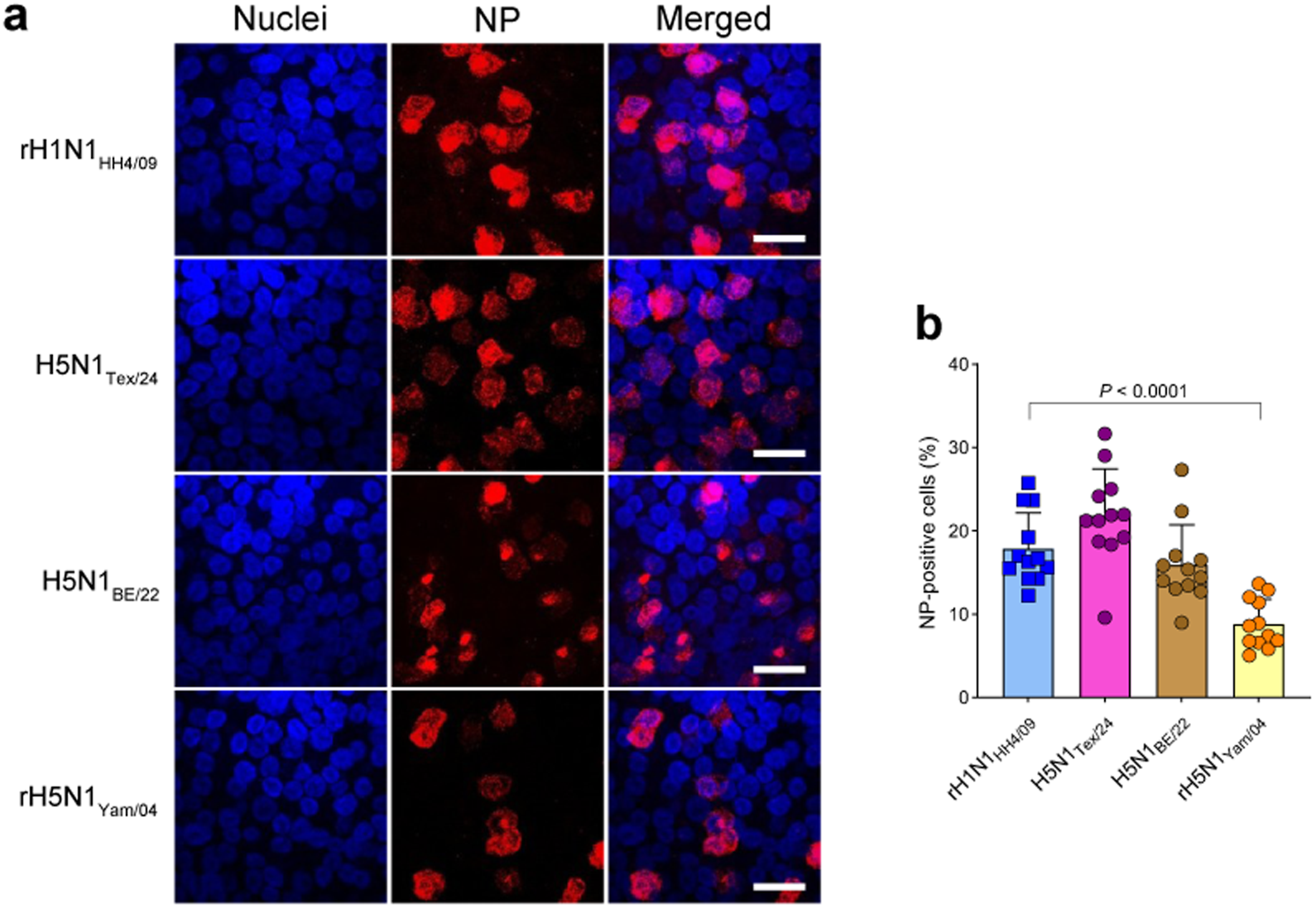
Infection efficiency of influenza viruses in HNE cells. **a, b** HNE cells differentiated at the air-liquid interface were infected with the indicated viruses at an MOI of 1 and maintained for 8 h at 37 °C. The cells were fixed with formalin and permeabilized with Triton X-100. Cells were stained for NP antigen and nuclei were stained by DAPI. Representative images from two infection experiments are shown. Bar size, 25 µm (**a**). The proportion of infected cells was determined as the percentage of nucleoprotein (NP)-positive cells relative to the total number of nuclei per field. Twelve fields per condition, each containing approximately 100 cells, were analyzed (**b**). Statistical significance was determined by one-way ANOVA with Dunnett′s multiple comparisons test. Only statistically significant *P* values are shown.

**Supplementary Fig. 2.**
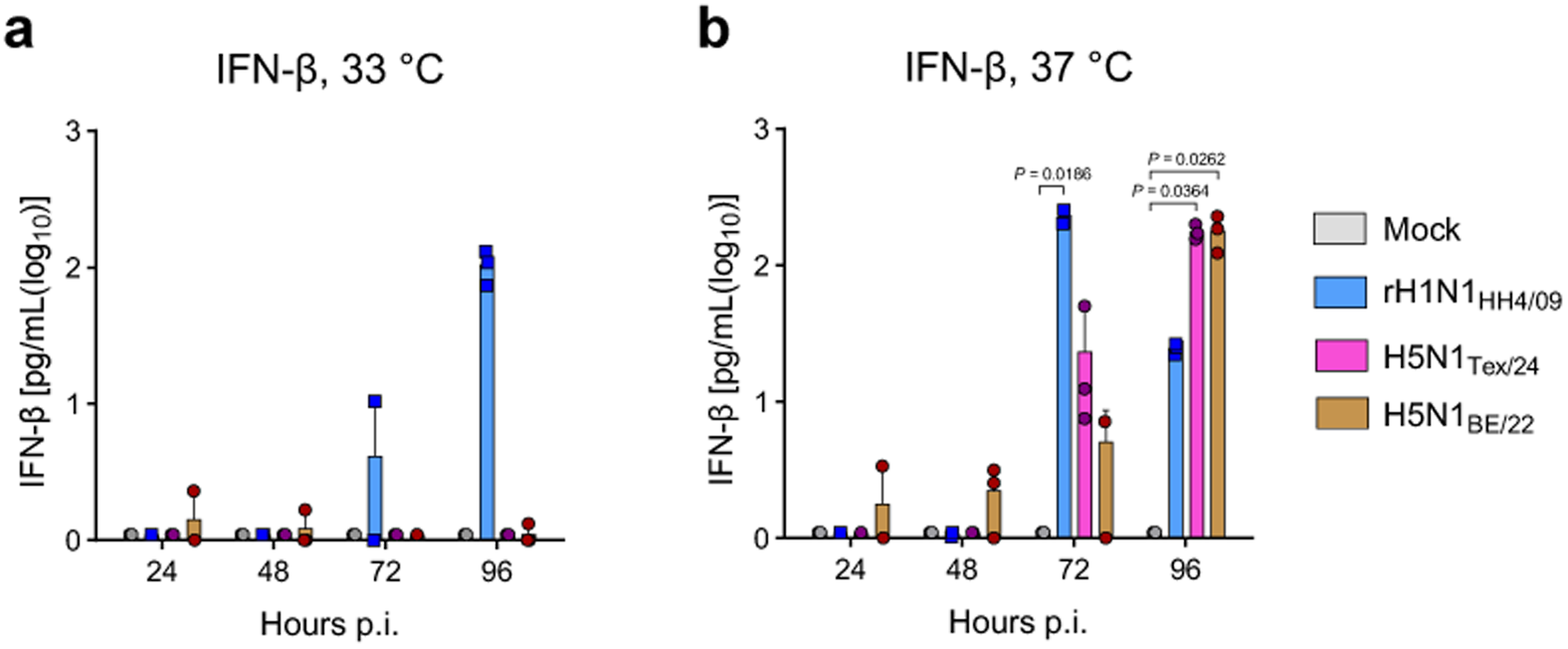
Detection of IFN-β in the basolateral medium of infected HNE cultures. **a, b** HNE cells differentiated at the air-liquid interface were infected with the indicated viruses at an MOI of 0.001 and incubated for 96 h at either 33 °C (**a**) or 37 °C (**b**). At the indicated time points, basolateral medium was collected and analyzed for IFN-β production by ELISA. Data represent mean values ± SD from three infection experiments. Statistical significance was determined by non-parametric Kruskal-Wallis with Dunn′s multiple comparisons test using mock-infected cells as reference. Only statistically significant *P* values are shown in the graphs.

**Supplementary Fig. 3.**
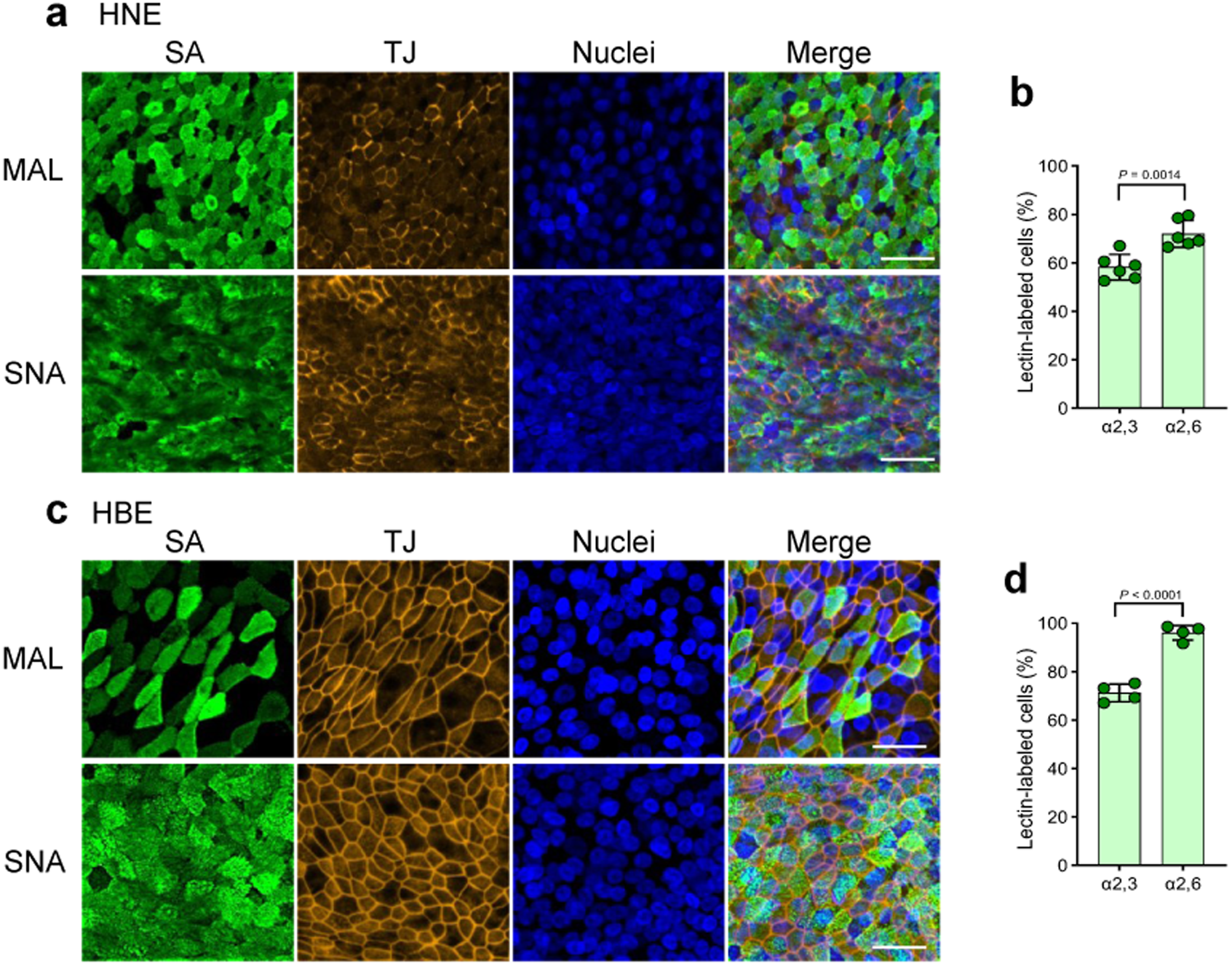
Detection of avian- and human-type receptors in HNE and HBE cells. **a - d** Human nasal epithelial (HNE) (**a, b**) and human bronchial epithelial (HBE) (**c, d**) cells were differentiated at the air-liquid interface at 33 °C and 37 °C, respectively. Cells were stained for the tight junction protein ZO-1 (TJ), α2,3-linked sialic acids (SA) using *Maackia amurensis* lectin (MAL), and α2,6-linked SA using *Sambucus nigra* agglutinin (SNA). Nuclei were counterstained with DAPI. Scale bar, 50 µm. The relative proportion of MAL- and SNA-positive HNE (**b**) and HBE (**d**) cells was quantified using ImageJ. Four to six images per condition, each containing approximately 100 cells, were analyzed. Data represent mean values ± SD. Statistical significance was determined using an unpaired two-tailed Student’s *t*-test.

