## Supplementary Information for "A bovine-derived influenza A virus (H5N1) shows efficient replication in well-differentiated human nasal epithelial cells at the temperature of the upper airways"

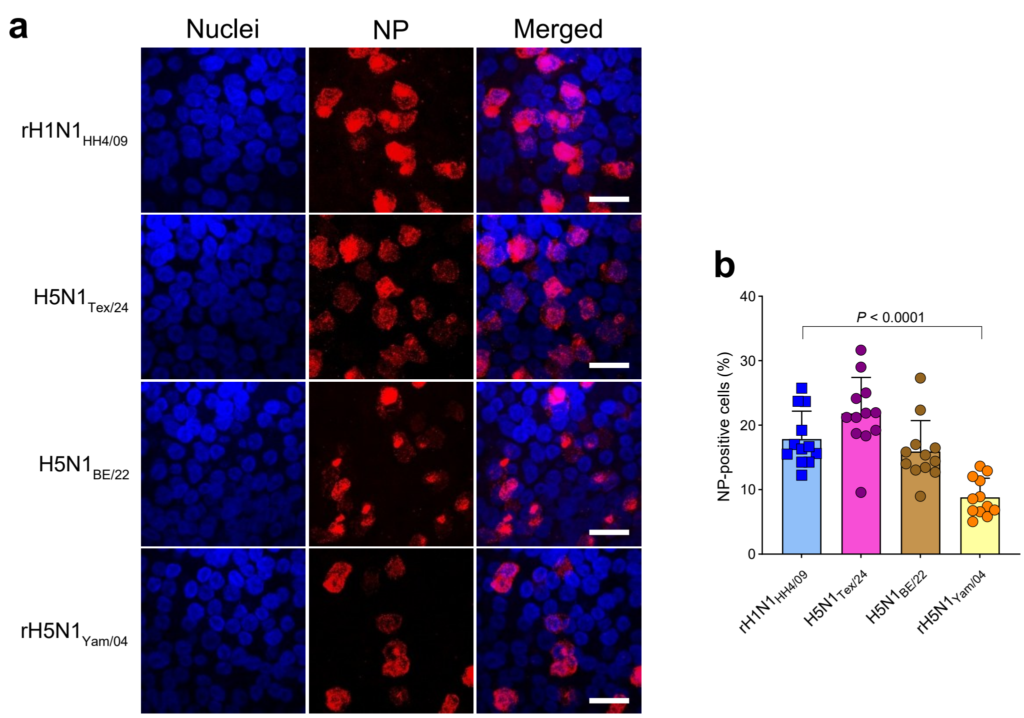


**Supplementary Fig. 1 | Infection efficiency of influenza viruses in HNE cells. a, b** HNE cells differentiated at the air-liquid interface were infected with the indicated viruses at an MOI of 1 and maintained for 8 h at 37 °C. The cells were fixed with formalin and permeabilized with Triton X-100. Cells were stained for NP antigen and nuclei were stained by DAPI. Representative images from two infection experiments are shown. Bar size, 25 µm (**a**). The proportion of infected cells was determined as the percentage of nucleoprotein (NP)-positive cells relative to the total number of nuclei per field. Twelve fields per condition, each containing approximately 100 cells, were analyzed (**b**). Statistical significance was determined by one-way ANOVA with Dunnett′s multiple comparisons test. Only statistically significant *P* values are shown.

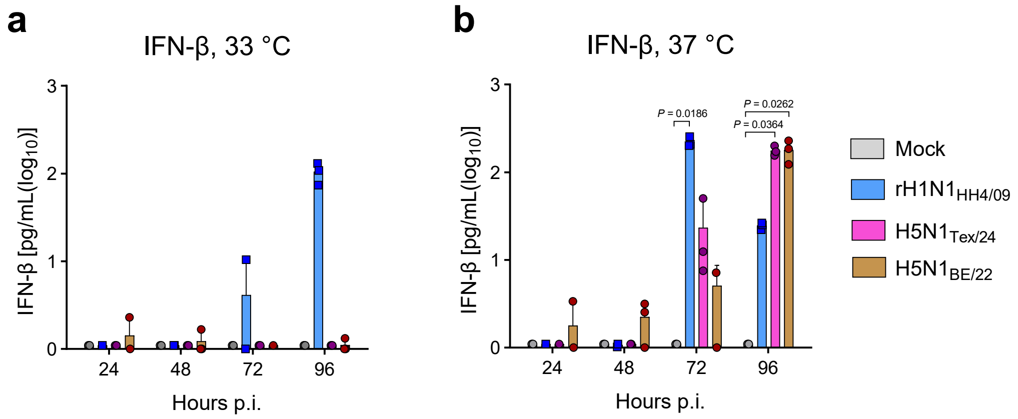


**Supplementary Fig. 2 | Detection of IFN-β in the basolateral medium of infected HNE cultures. a, b** HNE cells differentiated at the air-liquid interface were infected with the indicated viruses at an MOI of 0.001 and incubated for 96 h at either 33 °C (**a**) or 37 °C (**b**). At the indicated time points, basolateral medium was collected and analyzed for IFN-β production by ELISA. Data represent mean values ± SD from three infection experiments. Statistical significance was determined by non-parametric Kruskal-Wallis with Dunn′s multiple comparisons test using mock-infected cells as reference. Only statistically significant *P* values are shown in the graphs.

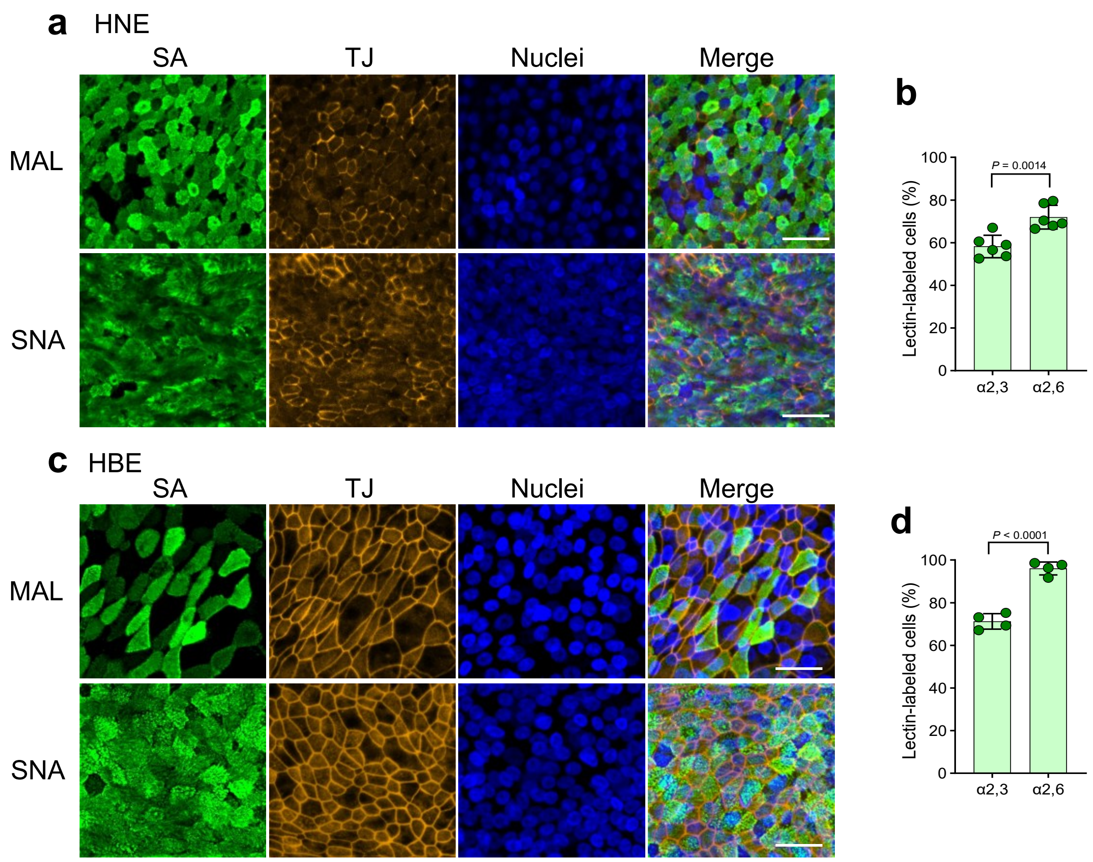


**Supplementary Fig. 3 | Detection of avian- and human-type receptors in HNE and HBE cells. a - d** Human nasal epithelial (HNE) (**a, b**) and human bronchial epithelial (HBE) (**c, d**) cells were differentiated at the air-liquid interface at 33 °C and 37 °C, respectively. Cells were stained for the tight junction protein ZO-1 (TJ), α2,3-linked sialic acids (SA) using *Maackia amurensis* lectin (MAL), and α2,6-linked SA using *Sambucus nigra* agglutinin (SNA). Nuclei were counterstained with DAPI. Scale bar, 50 µm. The relative proportion of MAL- and SNA-positive HNE (**b**) and HBE (**d**) cells was quantified using ImageJ. Four to six images per condition, each containing approximately 100 cells, were analyzed. Data represent mean values ± SD. Statistical significance was determined using an unpaired two-tailed Student's *t*-test.
